# CTCF and transcription influence chromatin structure re-configuration after mitosis

**DOI:** 10.1101/2021.06.27.450099

**Authors:** Haoyue Zhang, Jessica Lam, Di Zhang, Yemin Lan, Marit W. Vermunt, Cheryl A. Keller, Belinda Giardine, Ross C. Hardison, Gerd A. Blobel

**Affiliations:** Institute of Molecular Physiology, Shenzhen Bay Laboratory, Shenzhen, Guangdong, China; Division of Hematology, The Children’s Hospital of Philadelphia, Philadelphia, PA, USA; Perelman School of Medicine, University of Pennsylvania, Philadelphia, PA, USA; Department of Biochemistry and Molecular Biology, Pennsylvania State University, University Park, PA, USA

## Abstract

During mitosis, transcription is globally attenuated and chromatin architecture is dramatically reconfigured. Here we exploited the M- to G1-phase progression to interrogate the contributions of the architectural factor CTCF and the process of transcription to re-sculpting the genome in newborn nuclei. Depletion of CTCF specifically during the M- to G1-phase transition altered the re-establishment of local short-range compartmentalization after mitosis. Chromatin domain boundary reformation was impaired upon CTCF loss, but a subset (∼27%) of boundaries, characterized by transitions in chromatin states, was established normally. Without CTCF, structural loops failed to form, leading to illegitimate contacts between *cis*-regulatory elements (CREs). Transient CRE contacts that are normally resolved after telophase persisted deeply into G1-phase in CTCF depleted cells. CTCF loss-associated gains in transcription were often linked to increased, normally illegitimate enhancer-promoter contacts. In contrast, at genes whose expression declined upon CTCF loss, CTCF seems to function as a conventional transcription activator, independent of its architectural role. CTCF-anchored structural loops facilitated formation CRE loops nested within them, especially those involving weak CREs. Transcription inhibition did not elicit global architectural changes and left transcription start site-associated boundaries intact. However, ongoing transcription contributed considerably to the formation of gene domains, regions of enriched contacts spanning the length of gene bodies. Notably, gene domains formed rapidly in ana/telophase prior to the completion of the first round of transcription, suggesting that epigenetic features in gene bodies contribute to genome reconfiguration prior to transcription. The focus on the de novo formation of nuclear architecture during G1 entry yielded novel insights into how CTCF and transcription contribute to the dynamic re-configuration of chromatin architecture during the mitosis to G1 phase progression.

## Introductory paragraph

The mitotic phase of the cell cycle is characterized by rapid and extensive re-organization of chromatin architecture and global attenuation of transcription^1–5^. Studies of chromatin dynamics during entry into and exit from mitosis have informed the mechanistic basis underlying the hierarchical organization of chromatin. During mitotic exit, A/B compartmentalization is detectable as early as in ana/telophase but intensifies and expands thereafter^1–5^. Contacts between CREs such as promoters and enhancers are re-established with variable kinetics, some forming gradually and plateauing deeper into G1 while others are transient in nature being developed fully in ana/telophase only to fade upon G1-entry^1^. Resumption of transcription follows similarly variable characteristics: some genes display a spike in activity early in G1 and settle down at later stages whereas others are activated in a gradual fashion^1, 6, 7^.

The multi-functional transcription factor CTCF frequently co-localizes with boundaries of contact domains, such as topologically associated domains (TADs), and is proposed to assist their formation in collaboration with the cohesin ring complex through a process termed “loop extrusion” ^8–10^. Accordingly, acute depletion of CTCF or cohesin leads to wide-spread weakening of boundaries in interphase cells ^11–15^. CTCF and cohesin are evicted from mitotic chromatin to varying extents ^1, 16–19^, and measuring the rates by which they return to chromatin has enabled correlative assessments of their roles in post-mitotic genome folding and transcriptional activation. Upon mitotic exit, CTCF is immediately recruited back to chromatin prior to the formation of domain boundaries and architectural loops ^1^. The rate limiting step in the formation of these latter structures appears to be the accumulation of cohesin at CTCF bound sites, which occurs more gradually as chromatid extrusion proceeds.

Another feature frequently associated with domain boundaries is transcription start sites (TSS) ^20, 21^, but the role of transcription in boundary formation is still being debated. Inhibition of transcription compromises boundary strength in *Drosophila melanogaster* embryos^20, 21^. Yet, neither genetic nor chemical inhibition of transcription elicited a significant impact on higher-order structures of mammalian genomes ^22, 23^. A recent gain-of-function study demonstrated that ectopic insertions of TSSs can lead to the formation of new domain-like structures spanning the lengths of the de novo transcripts ^24^.

Most studies on how CTCF depletion or transcription inhibition impact chromatin architecture have been carried out in asynchronously growing cells and thus did not distinguish requirements for establishment versus maintenance of genome structure^11, 13–15, 20–22, 25^. The transition from mitosis into G1-phase offers the opportunity to monitor the *de novo* formation of compartments, compartment domains (here referring to genomic segments of a given compartment), domain boundaries, and chromatin loops in relation to CTCF binding and gene activation. Here, we interrogated the contributions of CTCF and the process of active transcription to the establishment of post-mitotic chromatin architecture by acutely depleting CTCF through the auxin inducible degron (AID) system alone or jointly with chemical inhibition of transcription during the mitosis to G1-phase transition ^1, 11, 26^.

## Results

### Cell cycle stage specific degradation of CTCF

To explore the impact of CTCF loss specifically during the period when chromatin architecture is rebuilt, we employed a murine erythroblast line G1E-ER4 in which both CTCF alleles were engineered to contain a C-terminal fusion to the AID-mCherry domains (Extended Data Fig. 1a) ^11, 26^. A TIR-expressing construct was transduced into the cells to allow for rapid auxin induced CTCF degradation (Extended Data Fig. 1a). CTCF became virtually undetectable after 1h exposure to auxin (Extended Data Fig. 1b). The acute nature of AID mediated degradation enabled removal of CTCF at precisely chosen time points. We applied auxin during nocodazole-induced prometaphase-arrest/release (Fig. 1a). CTCF depleted cells were enriched by FACS at defined time points during the prometaphase-to-G1 phase transition on the basis of mCherry fluorescence signal and DNA content staining (Extended Data Fig. 1c) ^1, 27^. To facilitate FACS purification of cells at ana/telophase, we adopted GFP fused to a mitotic-specific degron (MD) (Extended Data Fig. 1a). Purified cells were processed for *in-situ* Hi-C to detect architectural alterations (Extended Data Fig. 1d; Supplementary table 1). Short-term depletion of CTCF did not impede cell cycle progression, and accordingly, post-mitotic Hi-C contact decay curves were highly similar between auxin treated and control cells (Extended Data Fig. 1c; 2a), enabling pair-wise comparisons between CTCF depleted and replete cells at each post-mitotic cell cycle stage.

**Figure 1.**
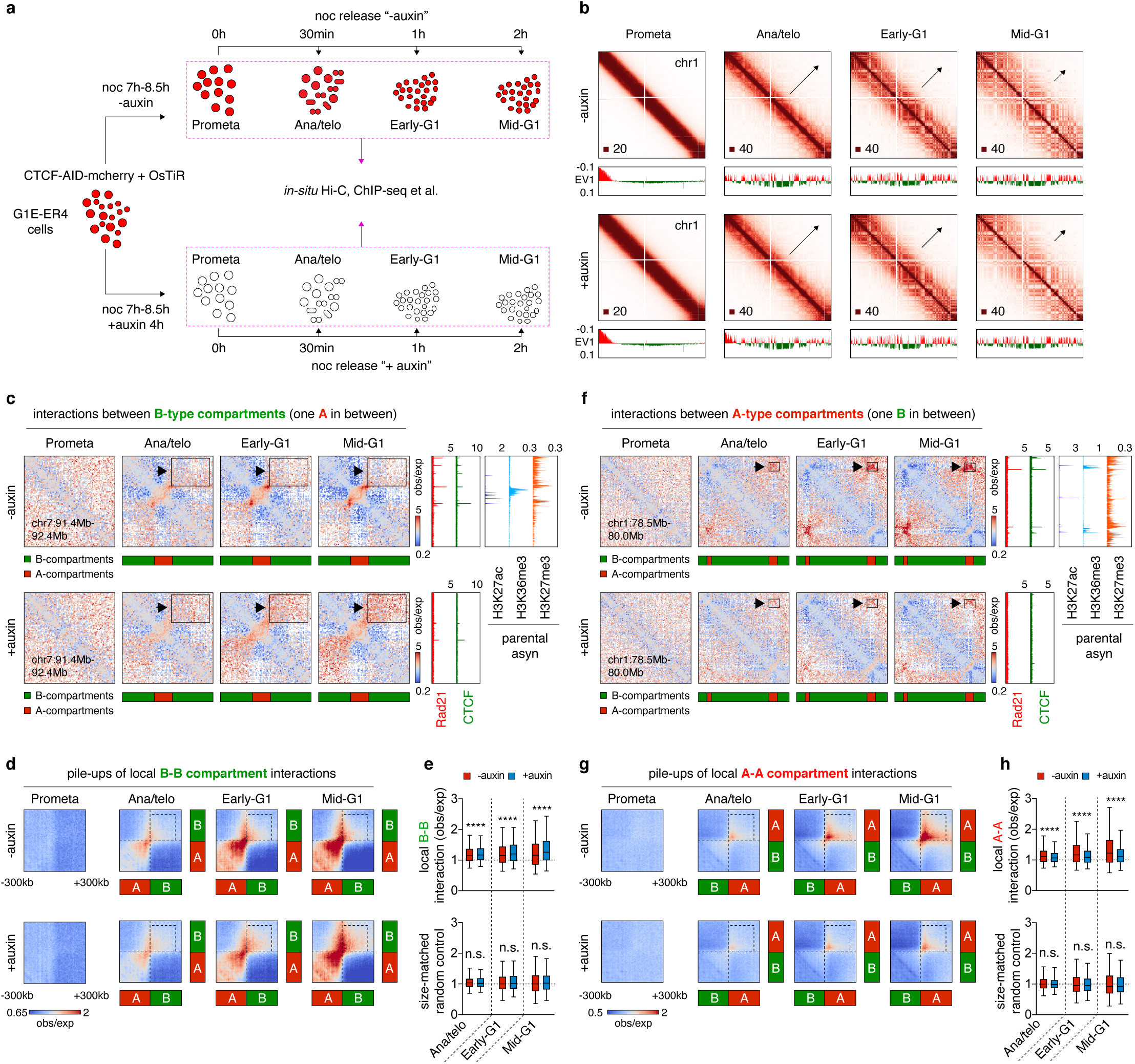
Alteration of local compartmentalization upon CTCF removal. **a**, Strategy for harvesting mitotic and post-mitotic populations with or without CTCF. **b**, KR balanced Hi-C contact matrices showing global compartment reformation of chr1 in untreated and auxin treated cells after mitosis. Bin size: 100kb. Black arrows indicate the progressive spreading of compartments throughout the entire chromosome. Browser tracks with compartment PC1 values are shown for each contact map. **c**, KR balanced Hi-C contact matrices showing representative local B-B interaction changes with or without CTCF depletion after mitosis. Bin size: 10kb. Arrows and boxes highlight the increased local B-B interactions after CTCF depletion across cell cycle stages. Tracks of CTCF and Rad21 with or without auxin treatment as well as histone marks H3K27ac, H3K36me3 and H27me3 are from asynchronous G1E-ER4 cells. **d**, Pile-up Hi-C matrices showing the increased local interactions between all consecutive (with one A-type in between) B-type compartment domains. Bin size: 10kb. Dotted boxes indicate the increased local B-B interactions genome-wide. **e**, Upper panel: Boxplots showing quantification of interactions in the dotted boxes (250kb x 250kb) in (**d**). Lower panel: Boxplots showing the effect of CTCF depletion on the interactions between randomly selected genomic pairs (n=500) that are distance-matched to the upper panel. For all boxplots, central lines denote medians; box limits denote 25th– 75th percentile; whiskers denote 5th–95th percentile. * *P* < 0.05, ** *P* < 0.01, *** *P* <0.001 and **** *P* <0.0001. Two-sided paired Wilcoxon signed-rank test. **f-h**, Similar to (**c-e**), showing examples, pile-ups and quantification of local consecutive (with one B-type in between) A-A interactions genome-wide.

### Local compartmentalization after mitosis requires CTCF

A/B compartment emergence in ana/telophase as well as expansion and intensity gains occurred at comparable rates in control and CTCF depleted cells, suggesting that CTCF is dispensable for global compartmentalization after mitosis (Fig. 1b; Extended Data Fig. 2b, d-g). Since prior reports suggested that loop extrusion counteracts compartmentalization ^12, 28^, we examined whether CTCF loss and the resulting extended extrusion process may affect short-range compartmental interactions. We analyzed the post-mitotic interaction frequency between consecutive B-type compartment domain pairs flanking a single A-type compartment domain and found it to be significantly increased upon CTCF depletion (Fig. 1c-e; Extended Data Fig. 3a). It is noteworthy that such increment of short-range B-B interactions grew as cells proceeded towards G1 (Fig. 1e; Extended Data Fig. 3h), in line with the progressive loading of cohesin after mitosis. Moreover, gains of B-B interactions only occurred locally and tapered off as they were further separated (Extended Data Fig. 3e-h), consistent with the limited residence time of cohesin on chromatin ^29^. Intriguingly, interactions between short-range A-type compartment domains were diminished in CTCF deficient cells (Fig. 1f-h; Extended Data Fig. 3b, e, g, i). Together, our data highlight a previously undescribed role of CTCF to regulate short-range compartmental interactions, likely by restricting cohesin driven loop extruding.

### CTCF dependent and independent mechanisms drive boundary reformation after mitosis

We next investigated post-mitotic boundary reformation upon CTCF depletion. We identified 6,376 boundaries with high concordance among biological replicates (Extended Data Fig. 4a, b; Supplementary table 2) ^30^. *K*-means clustering yielded five groups of boundaries with distinct sensitivities to CTCF depletion (Fig. 2a). Only ∼20% of boundaries (cluster1) were fully dependent on CTCF after mitosis (Fig. 2a; Extended Data Fig. 4c, f). ∼31.9% (cluster2) were partially dependent, and ∼27% (cluster3) were unaffected by CTCF loss (Fig. 2a; Extended Data Fig. 4d, e, g, h) indicative of CTCF independent mechanisms. As expected, cluster3 boundaries displayed markedly lower CTCF/cohesin occupancy than cluster 1 and 2 (Fig. 2b). Visual inspection suggested that cluster 2 and 3 boundaries are frequently located at transitions between regions enriched for the repressive histone mark H3K27me3 and the transcription elongation mark H3K36me3 (Extended Data Fig. 4i). We further quantified chromatin state transitions by principal component analysis (PCA) using 10kb binned H3K27me3 and H3K36me3 ChIP-seq signals across the -/+50kb region of each boundary (Fig. 2c). We found that the extreme (top or bottom 20%) PC1 projections reliably predicted transitions of chromatin states at boundaries (Fig. 2d). Importantly, cluster2 and 3 boundaries are significantly more enriched with top and bottom 20% PC1s compared to cluster1 (Fig. 2e). Therefore, cluster1 boundaries seem to be primarily driven by CTCF/cohesin mediated loop extrusion, Cluster3 boundaries may be formed through segregation of active and inactive chromatins, and cluster2 boundaries by both mechanisms. Intriguingly, under normal conditions (“-auxin”), cluster2 and 3 boundaries were reformed significantly faster than cluster1 after mitosis (Fig. 2f), suggesting that chromatin segregation may mediate more rapid insulation than loop extrusion after mitosis. Our data demonstrate that CTCF dependent and independent mechanisms can work separately or jointly to drive boundary formation after mitosis.

**Figure 2.**
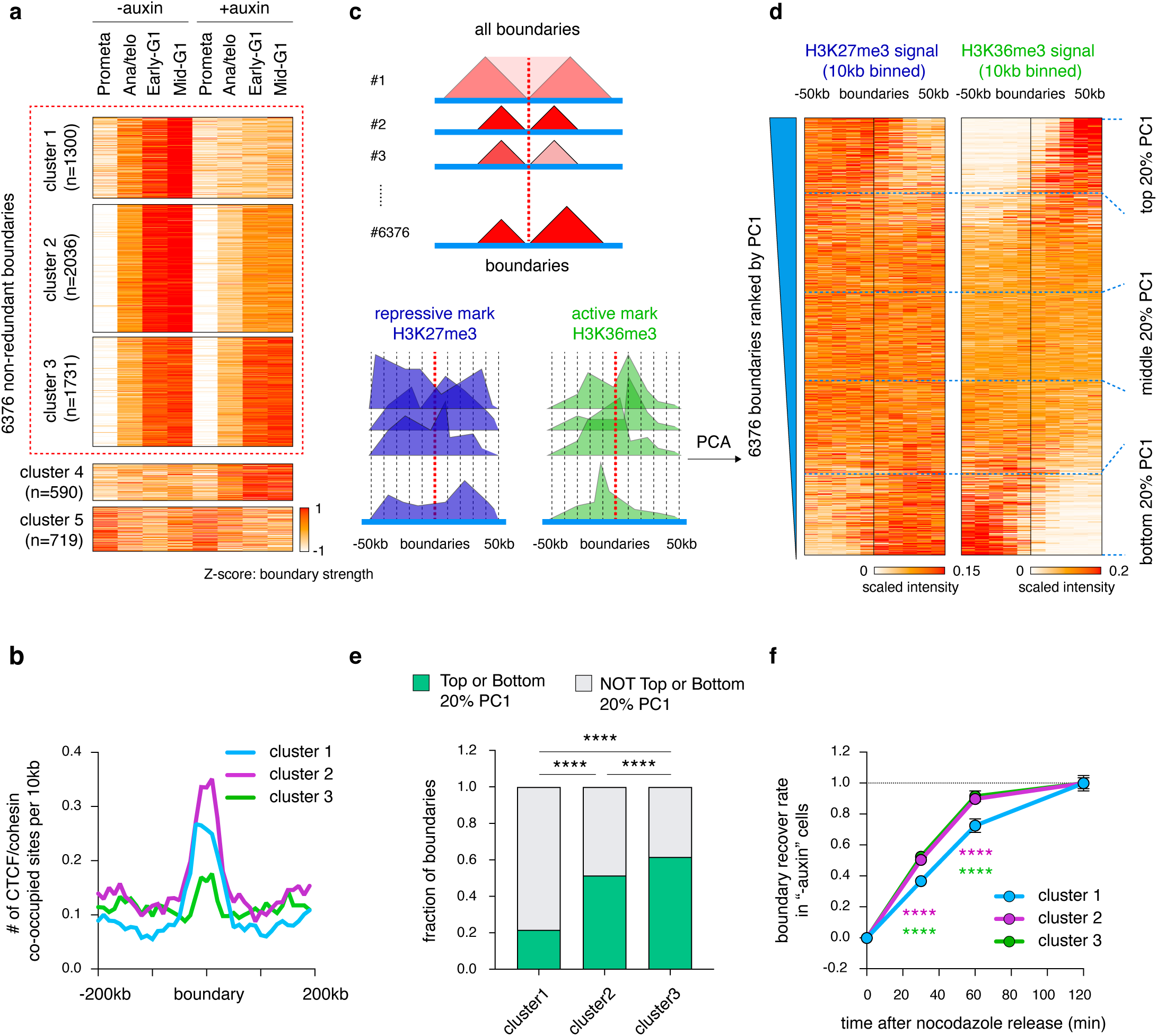
Reformation of boundaries display distinct responses to CTCF loss. **a**, *k*-means clustering of boundaries depending on their sensitivity to CTCF depletion. The z-scores in prometaphase for cluster1 boundaries should not be interpreted as the absolute insulation intensity, because they are calculated as relative values across all time points (see the absolute insulation intensity in Extended Data Fig. 4f-h). **b**, Average occupancy of CTCF/cohesin peaks per 10kb for boundaries from cluster 1-3. **c**, Schematic of the PCA based method using the H3k36me3 and H3K27me3 histone marks to assess boundaries as defined here as chromatin state transitions. **d**, ChIP-seq signal intensities of H3K27me3 and H3K36me3 in a 100kb window centered on boundaries. Boundaries were ranked by their PC1 projections in descending order. Top and bottom regions (20%) of the heatmap indicate transition of chromatin state from 5’ inactive to 3’ active and 5’ active to 3’ inactive respectively. **e**, Bar graphs showing the fraction of boundaries from each cluster with top or bottom 20% PC1 values. **** *P* <0.0001. *p* values were computed by Fisher’s exact test. **f**, Line graph showing the kinetics of boundary formation of clusters 1-3 in untreated cells. **** *P* <0.0001. Purple and green asteroids indicate *p* values from comparisons between cluster1 and cluster2 or 3 boundaries, respectively. Two-sided Mann-Whitney U test.

### Variable requirement of CTCF for post-mitotic loop formation

CTCF is frequently found at chromatin domain boundaries and anchors of architectural loops where it can promote or inhibit loop contacts among regulatory regions, such as enhancers and promoters. We stratified chromatin loops based on the composition of their loop anchors and asked to what extent they are rebuilt upon CTCF depletion. A modified HICCUPS algorithm ^1, 31^ identified a union of 16,370 loops across all time points and auxin treatment conditions with high concordance (Extended Data Fig. 5a, b; Supplementary table 3). Newly called loops were also appreciated visually at each cell cycle stage, supporting the validity of our loop calling method (Extended Data Fig. 5c). Among all loops, 8,207 (∼50%) harbor CTCF/cohesin co-occupied sites at both anchors. These were further sub-categorized into 4,837 “structural loops” with one or no anchor containing CREs (as defined previously^1^), and 3,370 “dual-function loops” with both anchors containing CREs (Fig. 3a). Post-mitotic reemergence of both structural and dual-function loops was severely disrupted upon CTCF loss as evidenced by aggregated peak analysis (APA) and PCA (Fig. 3b-d; Extended Data Fig. 5d). We also called 4,642 “CRE loops” with both anchors containing CREs and only one or no anchor harboring CTCF/cohesin peaks (Fig. 3a). While APA plots failed to reveal major changes in CRE loop establishment, ∼44.5% and ∼30.4% of CRE loops were lost and newly gained, respectively after auxin treatment (Fig. 3b, e, f), uncovering a considerable shift in their reformation after CTCF depletion. Our results thus point to a critical role for CTCF in the formation of diverse loop categories after mitosis.

**Figure 3.**
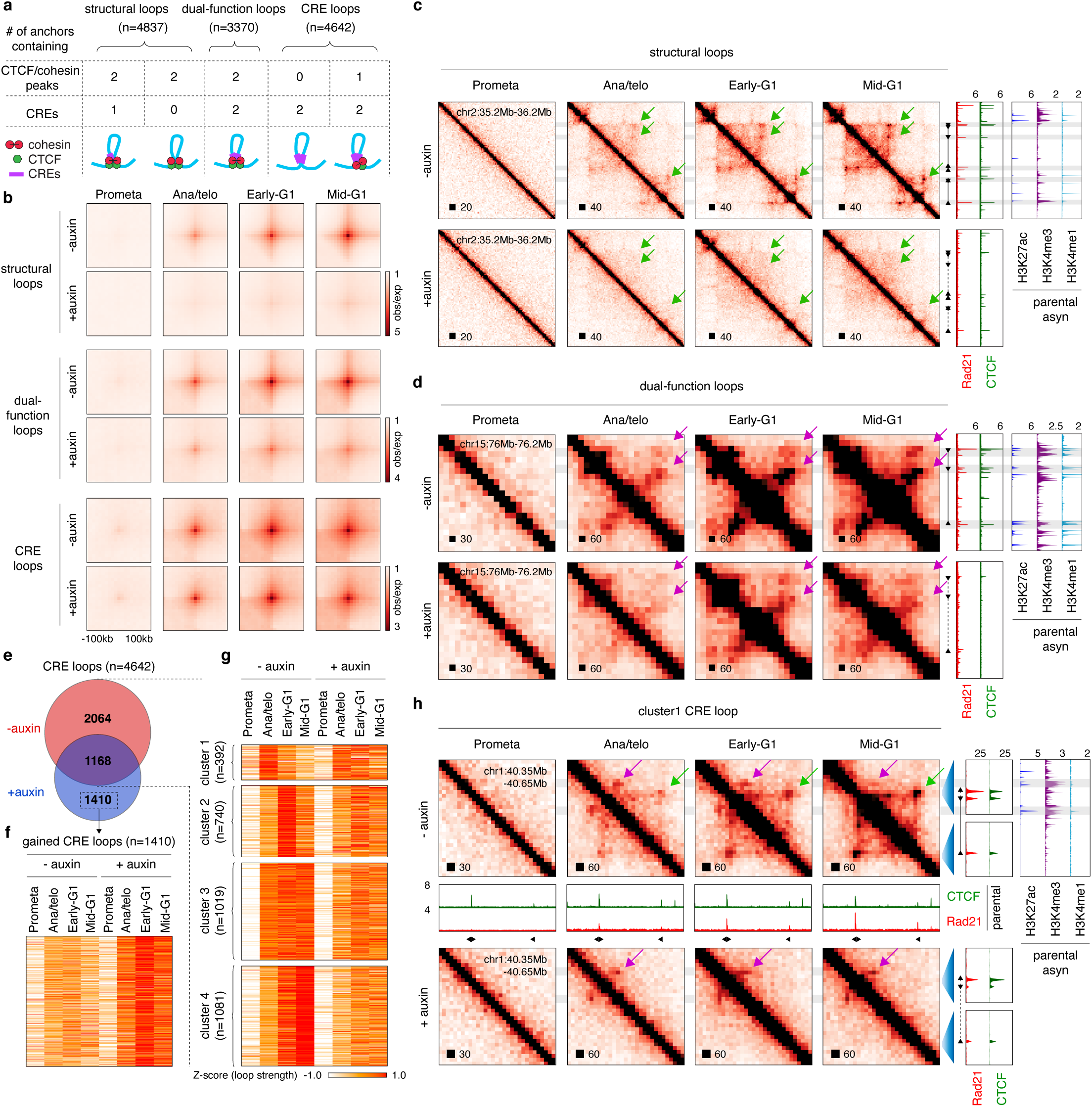
CTCF loops constrain CRE contacts after mitosis. **a**, Schematic showing the stratification of loops (“structural loops”, “dual-function loops” and “CRE loops”) based on the presence at their anchors of CTCF/cohesin co-occupied sites and CREs. **b,** APA plots showing the signals of loop categories before and after CTCF depletion across cell cycle stages. Bin size: 10kb. **c**, KR balanced Hi-C contact matrices of representative regions containing structural loops. Bin size: 10kb. Tracks of CTCF and Rad21 with or without auxin treatment as well as H3K27ac, H3K4me3 and H3K4me1 were from asynchronous G1E-ER4 cells. **d**, Similar to (**c**), KR balanced Hi-C contact matrices of representative regions containing dual-function loops. Bin size: 10kb. **e**, Venn diagram of CRE loops. **f**, Heatmap displaying intensities of the 1410 newly gained loops after CTCF depletion. **g**, Heatmap showing the result of *k*-means clustering on the 3232 CRE loops detected in untreated control samples. **h**, Similar to (**c**, **d**), KR balanced Hi-C contact matrices of a representative region containing a cluster1-P transient CRE loop. Additional tracks of CTCF and Rad21 from parental cells across designated cell cycle stages are shown^1^.

### Transient post-mitotic CRE loops are terminated by interfering structural loops

Previously, we uncovered a group of transient CRE loops whose intensities spiked in ana/telophase and subsequently faded in G1 ^1^. The function of such transient contacts if any, and the causative role for CTCF in their disruption remained unknown. Our acute CTCF depletion system enabled us to test this question globally. Using *k*-means clustering, we identified 392 transient CRE loops (cluster1) in control cells (Fig. 3g). Strikingly, 228 of these CRE loops persisted deep into G1 after CTCF depletion (cluster1-P), implying that CTCF is capable of blocking a subset of early-established CRE contacts after mitosis (Extended Data Fig. 6a, b). 164 CRE loops maintained their transient nature in the absence of CTCF (cluster1-NP) (Extended Data Fig. 6a, b). Visual examination of a representative cluster1-P CRE loop revealed the emergence of an interfering structural loop in control but not in auxin treated cells (Fig. 3h). Genome-wide, a high fraction (∼63.1%) of cluster1-P loops is potentially interrupted by structural loops, whereas remaining clusters (1-NP, 2, 3 & 4) were less affected (Extended Data Fig. 6c). Notably, the 1,410 newly gained CRE loops after CTCF loss also showed high level of structural loop interruption, comparable to that of cluster1-P (Extended Data Fig. 6c). This observation held true after randomized matching of CRE loop sizes from different clusters (Extended Data Fig. 6d).

CTCF resumes full chromatin occupancy in ana/telophase while cohesin accumulation is delayed^1^. Interference of CRE loops by structural loops occurs at later cell cycle stages (Fig 3h), suggesting that CTCF by itself is insufficient to block CRE interaction but requires engagement in cohesin mediated looped contacts. To test this idea, we performed genome-wide enrichment analysis to compare the likelihood of CRE loops to be disrupted by structural loops versus “loop-free” CTCF/cohesin co-occupied sites (Extended Data Fig. 6e). In comparison to structural loops, “loop-free” CTCF/cohesin sites displayed a reduced tendency to disrupt cluster1-P and the gained CRE contacts (Extended Data Fig. 6f). Furthermore, CTCF peaks independent of cohesin and structural loops were evenly distributed across all CRE loop clusters (Extended Data Fig. 6f). Together, dissection of the cell cycle dynamics of different loop categories uncovered support for the notion that CTCF is more effective as an insulator when part of a looped structure. A caveat of this interpretation is that small structural loops are undetectable in our Hi-C data.

Interposition of structural loops does not always disrupt CRE contacts: ∼37.5% of the CRE loops in clusters 2-4 had interposed structural loops. The failure of these to break up CRE loops might be related to their relative positioning. We observed that structural loops that are capable of weakening cluster1-P CRE loops tended to reside near (∼50kb) the CREs (Fig. 3h). To quantify this trait, we measured the distance of the influenced CRE to the most proximal structural loop anchor inside the CRE loop: s*_i_*-min (Extended Data Fig. 7a). Remarkably, s*_i_*-min was significantly shorter for cluster1-P and the gained CRE loops compared to the non-insulated cluster2, 3 and 4 (Extended Data Fig. 7b). This observation holds true for size-matched CRE loops (Extended Data Fig. 7c). Moreover, we observed a significant negative correlation between insulation strength and s*_i_*-min after mitosis (Extended Data Fig. 7d). Together, our data suggest that the insulating, CRE loop disrupting function of CTCF is linked not only to CTCF’s ability to form a loop but also to the relative position of the insulating loop.

### Structural loops can facilitate CRE connectivity after mitosis

CTCF depletion weakened a subset of CRE loops after mitosis (Fig. 3e), suggesting a supportive role of CTCF for certain CRE contact formation. APA plots demonstrated that the slower-forming cluster4 CRE loops were attenuated the most in G1 phase when compared to the other clusters (Extended Data Fig. 8a, b). Interestingly, cluster4 CRE loops were significantly more likely to reside within structural loops compared to size-matched ones from other clusters, implying a supportive role of structural loops for CRE-contacts (Extended Data Fig. 8c). Accordingly, the strengthening effects on cluster4 increased with cell cycle progression, consistent with the progressive re-formation of structural loops after mitosis (Extended Data Fig. 8d). It is noteworthy that anchors of cluster4 CRE loops displayed significantly weaker decoration with active histone marks H3K27ac, H3K4me1 and H3K4me3 than cluster3 (Extended Data Fig. 8e), suggesting that weak CRE contacts are especially reliant on encompassing structural loops.

### Reduced enhancer-promoter interactions do not account for transcription loss upon CTCF depletion

Given that CTCF loss altered post-mitotic CRE loop reformation, we examined how these changes impact gene reactivation after mitosis. We generated PolII ChIP-seq datasets during the mitosis-to-G1 phase transition with or without auxin treatment with high concordance among biological replicates (Extended Data Fig. 9a). We identified 7,238 active genes across all time points (Supplementary table 4). ∼52.0% of these genes showed post-mitotic transcriptional spiking in control cells (Extended Data Fig. 9b), as observed previously ^1, 6^. This spiking pattern was overwhelmingly maintained in the absence of CTCF (Extended Data Fig. 9b-d), suggesting that CTCF is not essential for re-activation of many genes, and validating that CTCF-deficient cells progress normally from prometaphase to G1. Consistent with previous reports, only small fraction of genes (426, ∼5.7%) were differentially expressed in at least one post-mitotic time point (*q* < 0.05, fold change > 1.25 fold) with 203 up-regulated and 223 down-regulated after mitosis (Fig. 4a) (see methods). Of note, most of these gene expression changes were already detectable in early-G1 phase, when transcription initiates, suggesting an instant effect of CTCF on these genes (Fig. 4a). The genes most down-regulated upon CTCF loss displayed the highest CTCF occupancy at their TSS in CTCF replete cells (Fig. 4b, c; Extended Data Fig. 9e, f), which could be explained by CTCF functioning as a direct transcription activator, or by mediating contacts with distal enhancers.

**Figure 4.**
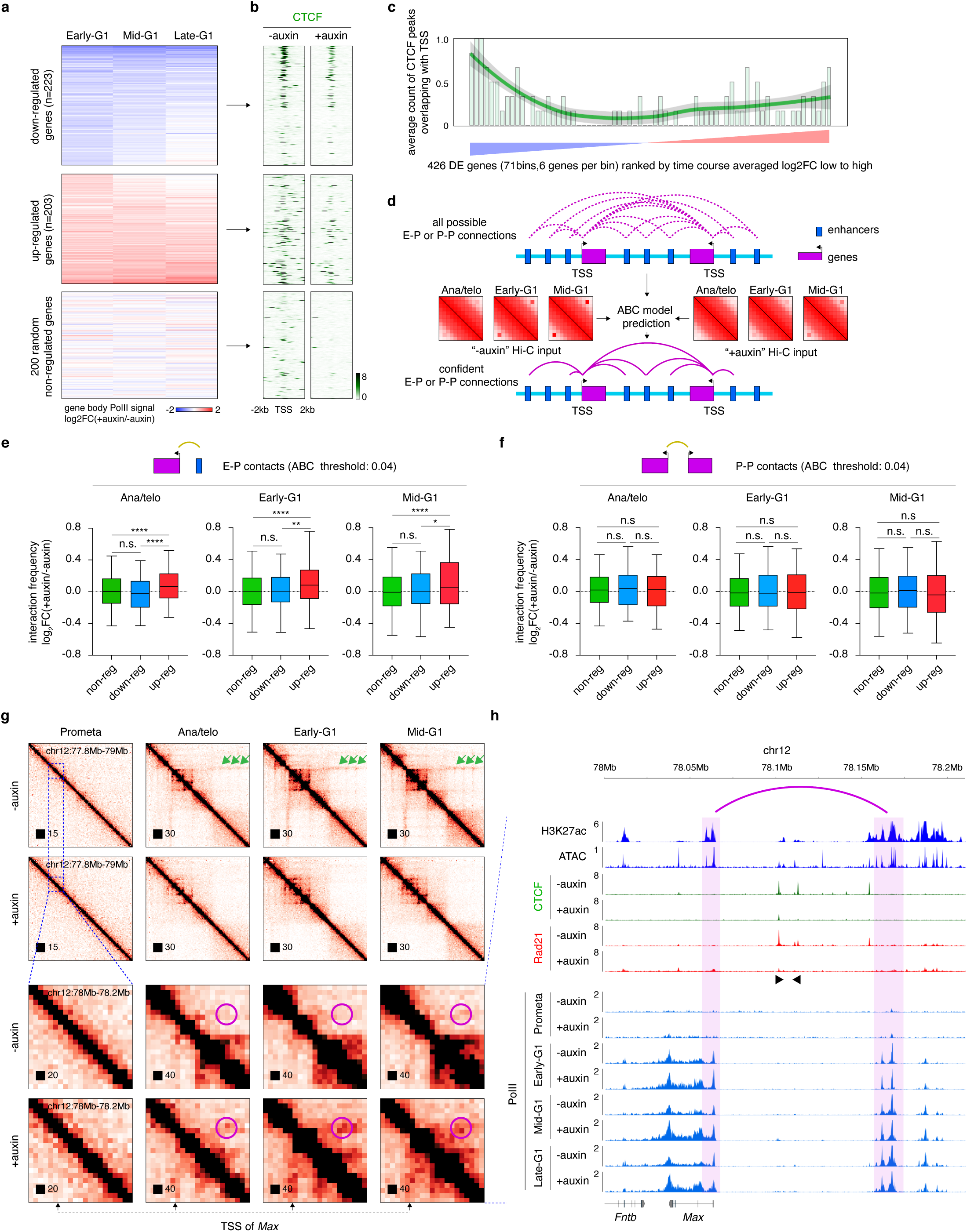
CTCF loss alters transcription reactivation profiles after mitosis. **a**, Heatmap displaying differentially expressed genes based on PolII ChIP-seq read counts over the gene bodies (+500 from TSS to TES), plotted as log2 fold-change (FC). **b**, Meta-region plots of CTCF ChIP-seq signals from asynchronous cells before and after auxin treatment, centered on down-regulated, up-regulated, or 200 random non-regulated gene TSS. **c**, Quantification of (**b**) showing the number of CTCF peaks overlaping with TSS. The green line represents lowess smoothing of bar plots. **d**, Schematic showing the implementation of the ABC model to predict confident E-P and P-P interactions using as input asynchronous H3K27ac ChIP-seq and ATAC-seq data from G1E-ER4 cells as well as *in-situ* Hi-C datasets from this study. **e**, Boxplots showing the log_2_ fold change upon CTCF depletion of interaction strength of E-P pairs (ABC score cutoff = 0.04) associated with either non-regulated, down-regulated, or up-regulated genes. log_2_ fold change of interaction strength was calculated using the LIMMA R package for each cell cycle stage. For all boxplots, central lines denote medians; box limits denote 25th–75th percentile; whiskers denote 5th–95th percentile. * *P* < 0.05, ** *P* < 0.01, *** *P* <0.001 and **** *P* <0.0001. Two-sided Mann-Whitney U test. **f**, Similar to (**e**) showing the interaction changes of promoter-promoter pairs after CTCF depletion. **g**, KR balanced Hi-C contact matrices showing the *Max* locus across cell cycle stages in control and auxin treated samples. Bin size: 10kb. Green arrows indicate the structural loops that insulate *Max* promoter from a nearby enhancer. Purple circles demarcate the increase of interactions between *Max* promoter and a nearby enhancer upon CTCF depletion after mitosis. Note that the gain in interactions occurs at the earliest tested time point. **h**, ChIP-seq genome browser tracks of the same region as that shown in the lower panel in (**g**). Note increased expression of *Max* after mitosis in auxin treated samples. Purple arch annotates the elevated interaction between *Max* promoter and nearby enhancer. Black arrows indicate the motif orientation of CTCF binding sites.

To distinguish between these possibilities, we implemented the activity-by-contact (ABC) model to call high confidence enhancer-promoter (E-P) contacts^32^ (Fig. 4d). Using a stringent ABC score threshold of 0.04 (see methods), we identified 7,725 E-P pairs associated with active genes. Unexpectedly, E-P pairs associated with down-regulated genes showed no significant reduction of contact intensity upon CTCF loss across all tested cell cycle stages compared to non-regulated genes (Fig. 4e; Extended Data Fig. 10a). This argues against CTCF-dependent looping as predominant mechanism to normally activate these genes, and suggests that CTCF functions as a transcriptional activator near the TSS.

### Gained enhancer-promoter interactions account for gene activation upon CTCF removal

We next explored how CTCF loss could lead to up-regulation of genes. We found that E-P pairs associated with up-regulated genes were significantly strengthened in the absence of CTCF after mitosis (Fig. 4e, g, h; Extended Data Fig. 10a). To quantify to what extent genes were regulated by distal enhancers, we set out to identify significantly altered E-P pairs (differential E-P interaction analysis) upon CTCF depletion. ∼20.2% of up-regulated genes (e.g. Max) displayed significantly strengthened E-P interactions after CTCF loss, while only ∼6% and ∼0.8% of non-regulated and down-regulated genes respectively were associated with increased E-P contacts (Extended Data Fig. 10b). This suggests that CTCF attenuates gene expression by interfering with E-P interactions. Accordingly, E-P pairs associated with up-regulated genes were more likely to be interrupted by structural loops, confirming an insulating role of structural loops in gene regulation (Extended Data Fig 10c). The above observations held across various ABC score thresholds (0.01-0.05), ruling out potential bias due to thresholding (Extended Data Fig. 10d, e). We also identified 11,766 promoter-promoter (P-P) pairs, which were essentially unchanged in all groups (up, down, non-reg) of genes (Fig. 4f; Extended Data Fig. 10a, e). This suggests that, in contrast to E-P contacts, P-P interactions contribute little to post-mitotic transcription reactivation. In sum, while overall CTCF depletion exerts modest effects on post-mitotic gene activation, proper regulation of some genes requires CTCF for their activation while others require it for shielding them from inappropriate enhancer influence.

### Compartment and boundary reformation was independent from transcription

The role of transcription in chromatin architecture is being debated. It was previously proposed that inhibition of transcription does not compromise boundary strength, while others reported that induction of transcription may lead to boundary formation at TSS and compartmental interactions changes ^25^. We next sought to investigate whether transcription facilitates post-mitotic compartment and boundary reformation. We treated cells with triptolide, a drug that inhibits transcription initiation, during the mitosis-to-G1 phase transition (Extended Data Fig. 11a) ^33, 34^, followed by *in-situ* Hi-C. Reformation of A/B compartments was ostensibly unperturbed in G1 upon transcription inhibition (Extended Data Fig. 2b-d, f). The CTCF independent cluster3 boundaries also remained intact after transcription inhibition (Extended Data Fig. 4h), ruling out active PolII complexes as underlying mechanisms. Moreover, insulation at TSS was unperturbed after transcription inhibition (Extended Data Fig. 11b). Consistent with this notion, insulation was progressively gained after mitosis at the TSS of the 100 most spiking genes even after their transcription was dialed down, suggesting that insulation as it occurs at TSS can be uncoupled from the process of transcription (Extended Data Fig. 11c-e). To examine whether transcription activity contributes to CTCF loss-induced chromatin changes, we removed CTCF and blocked transcription re-initiation simultaneously in mitosis and examined compartmentalization in G1 phase (Extended Data Fig. 11a). CTCF loss-induced alterations of local compartmental interactions (gain of B-B and loss of A-A) were also observed upon transcription inhibition (Extended Data Fig. 3c, d, j, k). CTCF loss-mediated increases in loop intensities for cluster1-P CRE loops were faithfully recapitulated in G1 upon transcription inhibition (Extended Data Fig. 6g). Furthermore, weakening of boundaries (cluster1 and 2 boundaries) as a result of CTCF withdrawal was recapitulated in triptolide treated cells (Extended Data Fig. 4f, g). Together, our results suggest that transcription is not a significant driving force for the formation of compartments and boundaries, or CTCF-dependent chromatin remodeling after mitosis.

### Transcription dependent and independent mechanisms drive gene domain formation at G1 entry

Active genes can appear as distinct squares on Hi-C contact maps, which has been interpreted to reflect gene domains caused by the process of transcription ^21, 22^. We examined to what extent transcription reactivation may dynamically influence genome reconfiguration by performing an integrative analysis of our Hi-C and post-mitotic PolII ChIP-seq datasets. A visually appreciable gene domain at the *Rfwd2* locus can be observed as early as in ana/telophase, even before PolII reached the transcription end site (TES) (Fig. 5a, b), suggesting that gene domains may appear quickly after mitosis prior to the completion of transcription elongation. To test this possibility genome-wide, we quantified the post-mitotic recovery rates of gene domains and gene-body PolII occupancy for all active genes. We found that small genes (30 to 50kb) displayed comparable rates of recovery between gene domains and PolII occupancy (Fig. 5c). However, the recovery rate of PolII was markedly reduced as gene size increased beyond 50kb and became significantly slower compared to that of gene domains (Fig. 5c, d; Extended Data Fig. 12a-g). Of note, CTCF removal had little impact on the recovery of gene domains after mitosis (Fig. 5c, d; Extended Data Fig. 12a-g). Transcription inhibition by triptolide diminished but did not abolish gene domain formation after mitosis (Extended Data Fig. 12h), indicating that active transcription accounts partially but not entirely for gene domain formation. Notably, gene domains were precisely decorated by H3K36me3. This mark delineates active gene bodies, but unlike the process of transcription itself, is stable throughout mitosis (Fig. 5b, d; Extended Data Fig. 12a-g)^34^. This suggests that re-establishment of gene domains that precedes the onset transcription might be facilitated by this chromatin mark, nominating H3K36me3 as potential mitotic bookmark.

**Figure 5.**
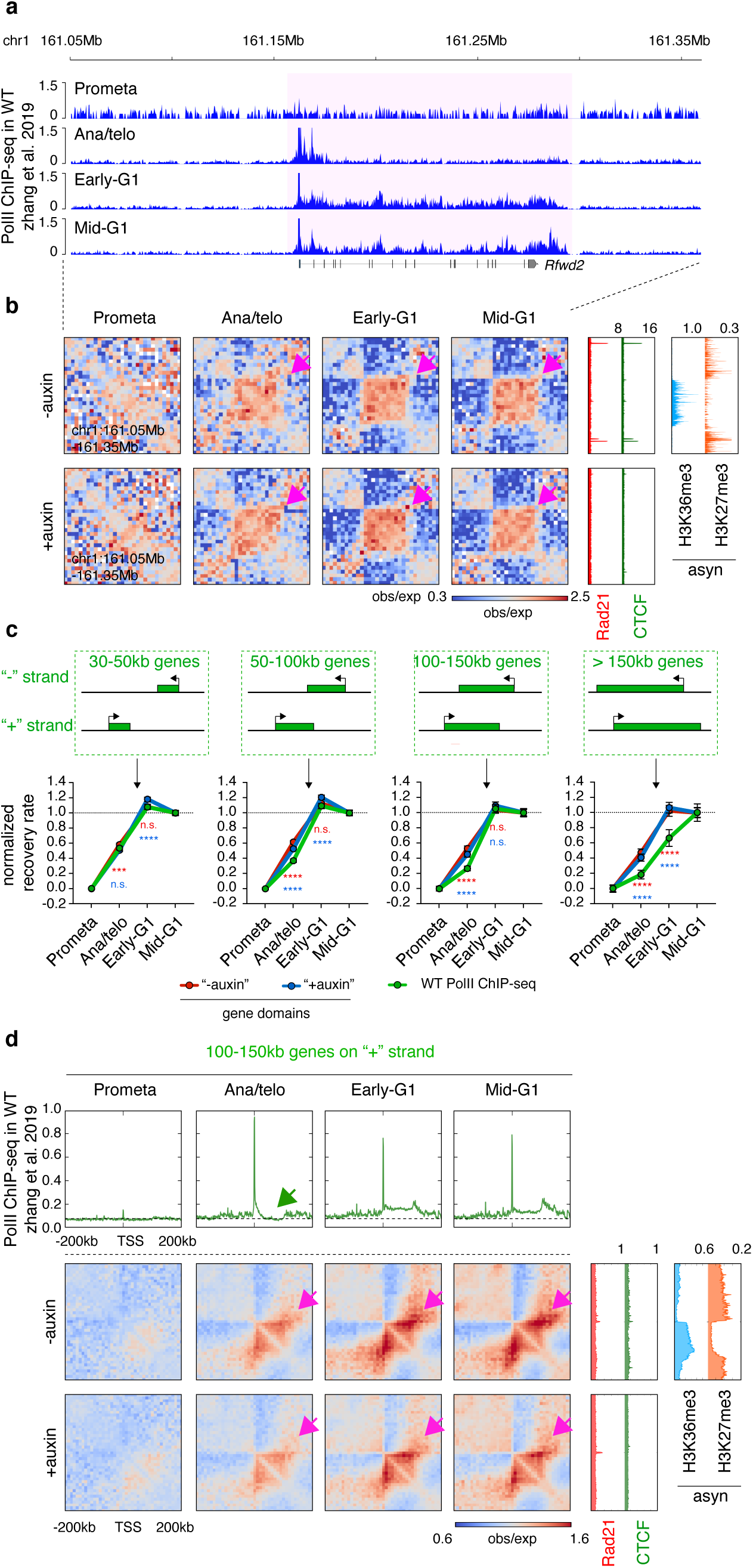
Gene domains emerge prior to completion of the first round of transcription after mitosis. **a**, PolII ChIP-seq genome browser tracks at the *Rfwd2* locus across cell cycle stages in parental cells. Note that in ana/telophase PolII is detected at the promoter region but the initial round of transcription has not been completed. **b**, KR balanced Hi-C contact matrices of the same region as in (**a**) across cell cycle stages in control and auxin treated samples. Bin size: 10kb. Purple arrows indicate domain of the *Rfwd2* gene in post-mitotic stages. Tracks of CTCF and Rad21 with or without auxin treatment as well as histone marks H3K36me3 and H27me3 are from asynchronously growing G1E-ER4 cells. **c**, Upper panel: Schematic of genes with different sizes. Lower panel: Line graphs of recovery rates of gene domains in the control and auxin treated samples, and the recovery rate of PolII occupancy over the gene body. Genes corresponding to the size ranges in the upper panel were separately plotted. *** *p* <0.001 and **** *p* <0.0001. Two-sided Mann-Whitney U test. Red and blue asterisks represent comparisons between PolII and gene domains in untreated control or auxin treated samples respectively. Error bars denotes SEM. **d**, Upper panel: Meta-region pile-up plots of PolII ChIP-seq signals corresponding to the 100kb-150kb genes on the plus strand across cell cycle stages. Plots are centered on TSS. Lower panel: Pile-up Hi-C matrices showing the domains of the genes corresponding to the upper panel across cell cycle stages in untreated and auxin treated samples. Bin size: 10kb. Plots are centered on TSS. Gene domains are labeled with purple arrows. Meta-region plots of CTCF and Rad21 with or without auxin treatment, as well as H3K36me3 and H3K27me3 are shown on the right.

## Discussion

Examination of the earliest stages of transition from pro-metaphase into G1 phase affords a unique view into how chromatin is configured de novo in newborn nuclei. The AID protein degradation system enabled investigation into the role of CTCF in this process. A meaningful interpretation of the experiments in this study requires that CTCF degradation does not significantly impede cell cycle progression. This was demonstrated by (1) flow cytometry measuring DNA content, cell size, and GFP-MD levels, (2) the presence of highly comparable contact decay curves across all post-mitotic time points, and (3) relatively stable post-mitotic gene expression patterns, including widespread gene spiking. As observed previously, only prolonged depletion of CTCF (36h) was found to delay cell cycle progression (Extended Data Fig. 1e), but this did not impact the present study.

Our data suggest that CTCF influences chromatin structure at several levels during G1 entry. First, CTCF based structural loops constrain short-range B-B compartmental interactions while promoting local A-A compartmental interactions, revealing a previously underappreciated role for CTCF in chromatin compartmentalization. We speculate that these observations are driven by altered loop extrusion after CTCF loss. Removal of CTCF may allow cohesin to travel beyond CTCF binding sites, thereby increasing loop sizes. Structural loops originating within A-type compartment domains may thus extend into flanking B-type compartment domains and increase their contact probability (Fig. 6a). The rising gains of B-B interactions during progression into G1 in the absence of CTCF are consistent with the gradual loading and advancement of cohesin after mitosis. The spatial confinement in B-B interaction gains might be due to limitations of the loop extrusion process. Our findings thus reveal that loop extrusion can elicit both positive and negative effects on compartmentalization depending on the type and location of the compartment (Fig. 6a)^12, 28, 35^. Second, the transient nature of many post-mitotic CRE contacts might be explained by the disruptive nature of emerging nearby structural loops. Thus, considering the cell cycle dynamics of structural as well as CRE loops during G1 progression allowed for the inference that CTCF’s ability to disrupt established CRE interactions, and hence function as an insulator, requires its engagement in loops (Fig. 6b). Third, we uncovered a previously underappreciated role of the genomic positioning of structural loop relative to CREs. Specifically, CRE contacts were most sensitive to disruption when the structural loop anchor was close to the CRE (small s*_i_*-min). While the mechanism underlying this observation is unclear, it is possible that the distance sensitivity of CRE contacts to the disruptive effects of extruding structural loops might be a function of the cohesin complex reaching and being arrested at CTCF sites more frequently. Fourth, CTCF-anchored loops may facilitate interactions between CREs by providing structural support. Weaker CREs appear to be more reliant on such “supportive” structural loops.

**Figure 6.**
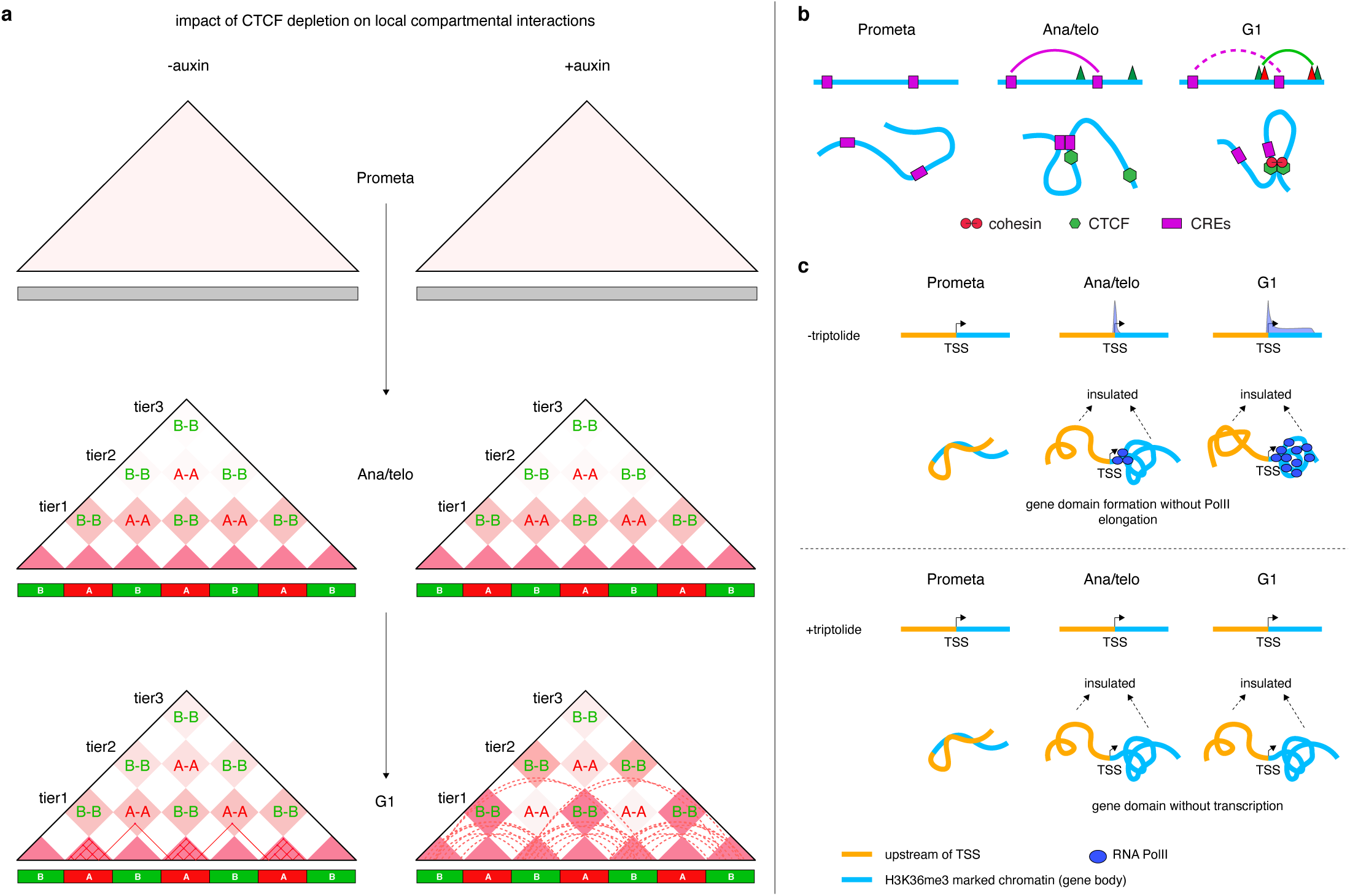
Mechanistic models. **a**, Schematic showing how CTCF removal can impact local but not distal interactions between same type of compartments. Short-range B-B interactions were enhanced potentially due to increased extrusion loop size from A compartments after CTCF removal. Note, the effect was progressively observed in G1 phase because of the gradual action of loop extrusion. Solid lines in the bottom panel represents structural loops formed and stabilized within A-type compartment domains in CTCF repleted conditions. Dotted lines represent actively extruding loops that are unleashed from A-type compartment domains into flanking B-type compartment domains due to CTCF depletion. **b**, Schematic showing the rapid dissolution of established CRE loops as nearby disruptive structural loop emerge after mitosis. **c**, Schematic showing that gene domain is well-established without full coverage of PolII over gene body. Effects after transcription inhibition was at the bottom panel.

Up-regulation of genes caused by CTCF loss was associated with enhanced interactions between promoters and enhancers, and was observed at the earliest measurements (1h after mitosis), suggesting a tight temporal relationship of promoter-enhancer proximity and transcription. Additionally, we found that the down-regulated genes generally did not display measurable loss in E-P contacts, but instead appear to depend on CTCF binding at their TSS. This implies that at these genes CTCF might function as a transcriptional activator, independent of its role in chromatin looping. This observation diverges from previous reports that CTCF depletion can diminish E-P interactions and result in transcription loss ^15, 36–38^. However, we cannot rule out that disruption of shorter range E-P loops, undetectable in our Hi-C experiments, might account for down-regulation gene transcription.

Prior studies reported a correlation between transcription activity and gene domains in asynchronously growing cells ^21, 22, 39^. However, comparing the kinetics of transcription re-activation and gene domain reformation, and inhibiting transcription pharmacologically revealed that the process of transcription *per se* does not account for the entirety of post mitotic gene domain formation (Fig. 6c). We speculate that additional mechanism (e.g. H3K36me3 histone modification along gene body) may contribute to self-aggregation of genes and pre-configure genes for subsequent activation.

In summary, by leveraging the auxin-inducible degron system and chemical transcription inhibition in the context of cell cycle dynamics, we were able to deepen our insights into the mechanisms by which CTCF and transcription reshape multiple facets of chromatin architecture from a randomly organized state in mitosis to the fully established structures in interphase.

## Acknowledgements.

We thank Andrew Katznelson and members of the Blobel lab for helpful discussions. We thank Dr. Elphege Nora from UCSF and Dr. Leonid Mirny from MIT for constructive comments. We thank the Flow cytometry core staff at the Children’s Hospital of Philadelphia for expert technical support on cell sorting. This work was supported by NIH grants DK058044 (G.A.B.) and R24DK106766 (R.C.H and G.A.B.).

## Author Contributions

H.Z. and G.A.B conceived the study and designed experiments. H.Z. performed all experiments with help from J.L., D.Z., A.K., M.V., C.K., B.G. and R.C.H. H.Z. performed all data analysis with help from Y.L. H.Z. and G.A.B wrote the paper with contribution from all listed authors.

## Materials and methods

### Cell culture and maintenance

G1E-ER4 cells were cultured in suspension as previously described and maintained at a density of not exceeding one million/μl ^40^. Construction of the G1E-ER4 sub-line with AID-mCherry tagged CTCF has been described previously ^1^. We expressed OsTiR-IRES-GFP (to isolate prometa, early-G1 and mid-G1 phase cells) or OsTiR-IRES-GFP-MD (to isolate ana/telophase cells) in G1E-ER4 CTCF-AID-mCherry cells with the retro-viral vector MigR1. OsTiR positive cells were enriched via FACS based on GFP signal.

To measure cell growth of G1E-ER4 CTCF-AID-mCherry cells with or without OsTiR-IRES-GFP, 10^5^ cells from each line were treated with or without 1mM auxin and cells were counted at 12h, 24h and 36h respectively.

### Validation of CTCF depletion upon auxin treatment

G1E-ER4 CTCF-AID-mCherry cells expressing OsTiR-IRES-GFP were treated with 1mM auxin for 0min, 30min, 60min, 120min or 240min. Cells were fixed with 1% formaldehyde and subject to flow cytometry for mCherry signal. Wildtype G1E-ER4 cells were used as control.

### Cell synchronization and auxin treatment

For “+auxin” samples:

To enrich cells at prometaphase, early-G1 phase or mid-G1 phase, the G1E-ER4 CTCF-AID-mCherry cells over expressing OsTiR-IRES-GFP were treated with nocodazole (200ng/ml) for 7-8.5h at a density of around 0.7-1million/ml. To degrade CTCF during mitosis, auxin (1mM) was added to the culture during the last 4 hours of nocodazole treatment so that CTCF was removed by the end of prometaphase synchronization. To acquire post-mitotic populations, nocodazole treated cells were pelleted at 1200rpm for 3min. Cells were then washed once and immediately re-suspended in warm nocodazole-free medium containing 1mM auxin for 60min (early-G1) and 120min (mid-G1) respectively. To enrich for cells at ana/telophase, G1E-ER4 CTCF-AID-mCherry cells over expressing OsTiR-IRES-GFP-MD were first synchronized as above described and then released from nocodazole for 30min before harvest.

Control samples underwent the exact same treatment except that no auxin was added.

### Transcription inhibition

For “+auxin” samples:

G1E-ER4 CTCF-AID-mCherry cells expressing OsTiR-IRES-GFP were arrested in prometaphase with nocodazole (200ng/ml) as above. Auxin was added during the last 4h of nocodazole treatment. Triptolide (1μM) was added to the cultures during the last hour of nocodazole exposure. Cells were released into warm nocodazole free medium with 1μM triptolide and with or without 1mM auxin for 2h.

### Cell sorting

Control and “+auxin” samples were acquired as described previously^1^. Briefly, for *in-situ* Hi-C experiments cells were pelleted at 1200rpm for 3min. Cells were then re-suspended in 1 x PBS and crosslinked with 2% formaldehyde for 10min at RT. Crosslinking was quenched with 1M (final concentration) glycine for 5min at RT. Cells were permeabilized by 0.1% Triton X-100 for 5min at RT and stained with antibody against the mitosis specific antigen pMPM2 (Millipore, 05-368, 0.2µl/10million cells) for 50min at RT. Cells were then treated with APC-conjugated F(ab’)2-goat anti-mouse secondary antibody (Thermo Fisher Scientific, 17-4010-82, 2µl/10million cells) for 30min at RT. Cells were pelleted and re-suspended in 1 x FACS sorting buffer (1 x PBS, 2% FBS, 2mM EDTA and 0.02% NaN_3_) containing 20ng/ml DAPI at a density of about 50-100 million cells/ml. Prometaphase cells were purified via FACS on the basis of pMPM2 signal (+) and DAPI signal (4N). To harvest populations after mitotic exit, cells were pelleted and crosslinked with 2% formaldehyde at designated time points (ana/telophase, early-G1 and mid-G1 phase). Crosslinking was halted with 1M (final concentration) glycine at RT, and cells were permeabilized with 0.1% Triton X-100. Finally, cells were re-suspended in 1 x FACS sorting buffer and subjected to FACS sorting. Ana/telophase cells were sorted based on GFP signal (reduced) and DAPI signal (4N). Early-G1 and mid-G1 cells were sorted based on DAPI signal (2N). In addition, mCherry positive and negative populations were gated to collect untreated control and auxin treated samples respectively for all mitotic and post-mitotic time points. Sorted cells were snap-frozen and stored at −80℃.

The exact same procedure was carried out for cells that had undergone triptolide treatment.

For PolII ChIP-seq: Cells with or without auxin treatment were harvested at 0min (prometa), 60min (early-G1), 120min (mid-G1) or 240min (late-G1) after nocodazole release. Cells were re-suspended in 1 x PBS and crosslinked with 1% formaldehyde for 10min at RT. Crosslinking was stopped with 1M glycine followed by permeabilization with Triton X-100. All samples (+/-auxin at all cell cycle stages) were stained with anti-pMPM2 antibody (Millipore, 05-368, 0.2µl/10million cells) for 50min at RT, followed by APC-conjugated F(ab’)2-goat anti-mouse secondary antibody (Thermo Fisher Scientific, 17-4010-82, 2µl/10million cells) for 30min at RT. Cells were re-suspended in 1 x FACS buffer containing DAPI and subjected to FACS sorting.

Prometaphase cells were purified via FACS based on pMPM2 signal (+) and DAPI signal (4N). Early-, mid- and late-G1 cells were sorted based on DAPI signal (2N). mCherry positive and negative populations were gated to collect untreated control and auxin treated samples for all mitotic and post-mitotic time points. Sorted cells were snap-frozen and stored at −80℃. Note that protease inhibitor and PMSF were added to all buffers during the entire sample preparation procedure.

### In-situ Hi-C

*In-situ* Hi-C experiments were performed as previously described ^1^. Briefly, sorted cells (5 million for prometaphase and ana/telophase and 10million for early- and mid-G1 phase) were lysed in 1ml cold Cell Lysis Buffer (10mM Tris pH 8, 10mM NaCl, 0.2% NP-40/Igepal) for 10min on ice. Nuclei were pelleted at 4℃ and washed with 1.2 x DpnII buffer. Nuclei were permeabilized with 0.3% SDS for 1h at 37℃ and quenched with 1.8% Triton X-100 for 1h at 37℃. Chromatin was digested with 300U DpnII restriction enzyme (NEB, R0543M) *in-situ* at 37℃ over night with shaking. 300U DpnII restriction enzyme was added for an additional 4h at 37℃ with shaking. Nuclei were incubated at 65℃ for 20min to inactivate DpnII. After cool down, digested chromatin fragments were blunted with pCTP, pGTP, pTTP and Biotin-14-dATP (Thermal Fisher Scientific, 19524016) using 40U DNA Polymerase I, Large (Klenow) fragment (NEB, M0210). DNA was ligated *in-situ* with 4000U T4 DNA ligase (NEB, M0202M) for 4h at 16℃ followed by further incubation for 2h at RT. Nuclei were then incubated in 10% SDS containing proteinase K (3115879 BMB) at 65℃ overnight to reverse crosslinking. RNA was then digested with DNase-free RNase at 37℃ for 30min. DNA was then extracted by phenol-chloroform extraction, precipitated, and dissolved in nuclease free water. DNA was sonicated to 200-300bp fragments (Epishear, Active Motif, 100% amplitude, 30s ON and 30s OFF, 25-30min) and purified with AMPure XP beads (Beckman Coulter). Biotin-labeled DNA was purified by incubation with 100µl Dynabeads MyOne Streptavidin C1 beads (Thermal Fisher Scientific, 65002) at RT for 15min. DNA libraries were constructed using the NEBNext DNA Library Prep Master Mix Set for Illumina (NEB E6040, M0543L, E7335S). To elute DNA, streptavidin bead-bound DNA was incubated in 0.1% SDS at 98℃ for 10min. DNA was purified with AMPure XP beads and index labeled with NEBNext multiplex oligos for 6 cycles on a thermal cycler, using the NEBNext Q5 Hot Start HIFI PCR master mix. Index labeled PCR products were then purified with AMPure XP beads and sequenced on an Illumina NextSeq 500 sequencer.

### ChIP-seq

Chromatin immunoprecipitation (ChIP) was performed using anti-RNA Polymerase II antibody (Cell Signaling, 14958) as described previously ^1^. Briefly, following sorting, cells were re-suspended in 1ml pre-cooled Cell Lysis Buffer supplemented with protease inhibitors (PI) and PMSF for 20min on ice. Nuclei were pelleted and re-suspended in 1ml Nuclear Lysis Buffer (50mM Tris pH 8, 10mM EDTA, 1% SDS, fresh supplemented with PI and PMSF) for 10min on ice. 0.6ml IP dilution buffer (20mM Tris pH 8, 2mM EDTA, 150mM NaCl, 1% Triton X-100, 0.01% SDS, fresh supplemented with PI and PMSF) was added followed by sonication (Epishear, Active Motif, 100% amplitude, 30s ON and 30s OFF) for 45min. Samples were pelleted at 15000rpm for 10min at 4℃ to remove cell debris. Supernatant was supplemented with 3.4ml IP dilution buffer fresh supplemented with PI and PMSF, 50μg isotope-matched IgG and 50μl protein A/G agarose beads (A:G = 1:1, ThermoFisher 15918014 and ThermoFisher 15920010) and rotated at 4℃ for 8h to preclear the chromatin. 200µl, chromatin was set aside as input chromatin. Precleared chromatin was then incubated with 35µl A/G agarose beads (A:G = 1:1) pre-bound with anti-RNA PolII antibody (5µg/IP) at 4℃ for overnight. Beads were washed once with IP wash buffer I (20mM Tris pH 8, 2mM EDTA, 50mMNaCl, 1% Triton X-100, 0.1% SDS), twice with high salt buffer (20mMTris pH 8, 2mM EDTA, 500mMNaCl, 1% Triton X-100, 0.01% SDS), once with IP wash buffer II (10mMTris pH 8, 1mM EDTA, 0.25 M LiCl, 1% NP-40/Igepal, 1% sodium deoxycholate) and twice with TE buffer (10mM Tris pH 8, 1mM EDTA pH 8). All washing steps were performed on ice. Beads were moved to RT and eluted in 200µl fresh made Elution Buffer (100mM NaHCO3, 1%SDS). 12µl of 5M NaCl, 2µl RNaseA (10mg/ml) was added to IP and input samples and incubated at 65℃ for 2h, followed by addition of 3µl protease K (20mg/ml) and incubated at 65℃ overnight to reverse crosslinking. Finally, IP and input samples were supplemented with 10ul of 3M sodium acetate (pH 5.2), and DNA was purified with QIAquick PCR purification kit (QIAGEN 28106). ChIP-seq libraries were constructed using the Illumina’s TruSeq ChIP sample preparation kit (Illumina, catalog no. IP-202-1012). Libraries were size selected using the SPRIselect beads (Beckman Coulter, catalog no. B23318) before PCR amplification. Libraries were then quantified through real-time PCR with the KAPA Library Quant Kit for Illumina (KAPA Biosystems catalog no. KK4835). Finally, libraries were pooled and sequenced on a Illumina NextSeq 500 platform using Illumina sequencing reagents.

### Quantification and data analysis

#### Hi-C data pre-processing

For each biological replicate, paired end reads were aligned to the mouse reference genome mm9 using bowtie2 (global parameters: --very-sensitive -L 30 --score-min L,-0.6,-0.2 --end-to-end –reorder; local parameters: --very-sensitive -L 20 --score-min L,-0.6,-0.2 --end-to-end --reorder) through the Hi-C Pro software ^41^. PCR duplicates were removed and uniquely mapped reads were paired to generate a validPair file. The output validPair file was converted into “.hic” file using the hicpro2juicebox utility. For merged samples, similar steps were taken on reads merged from each biological replicate.

#### A/B compartment calling and processing

Compartments were called based on the “.hic” files through eigenvector decomposition on the Pearson’s correlation matrix of the observed/expected value of 100kb binned, Knight-Ruiz (KR) balanced cis-interaction maps (Eigenvector utility of juicer_tools_1.13.02)^31^. Positive and negative EV1 values of each 100kb bin were assigned to A-(active) and B-(inactive) compartments, respectively, based on gene density. Compartments were called on both replicate-merged samples and individual biological replicates across all conditions. Chromosome 3 was excluded from compartment analysis due to a chromosomal translocation.

#### Saddle plotting and global compartment strength calculation

To visualize compartment strength, we generated saddle plots. Briefly, 100kb binned Knight-Ruiz (KR) balanced *cis* observed/expected contact matrix was extracted from each “.hic” file through the DUMP utility of juicer_tools (1.13.02)^31^. For untreated control samples, the contact matrices were transformed in the same way such that each row and column of bins was reordered based on the eigenvector 1 (EV1) values associated with the mid-G1 sample, so that they fall into an ascending order from top to bottom and from left to right. Similar transformation was applied onto auxin treated samples across all cell cycle stages, based on their mid-G1 samples. Also, similar transformation was performed on triptolide treated G1-phase samples either with or without auxin treatment. After the transformation, bins at the top-left corner are associated with B-B compartment interactions. Bins at the bottom-right corner are associated with A-A compartment interactions. Bins at the top-right and bottom-left corners are associated with B-A and A-B compartment interactions respectively. The transformed contact maps from each chromosome were divided into 50 equal sections and averaged to create the genome wide saddle plots. The compartment strength of each individual chromosome was computed as following: compartment strength = (median (top20% AA) + median (top20% BB)) / (median (top20% AB) + median (top20% BA)). The compartment strength from each individual chromosome was averaged and log2 transformed as genome wide compartment strength. The compartment strength of individual replicates and merged samples were computed independently.

#### Compartmentalization expansion curve *R(s)*

The analysis of the progressive compartmentalization spreading across cell cycle stages has been described previously^1^. *R(s)* curve was established to indicate the distance-dependent level of compartmentalization. To compute the *R(s)* curve, 100kb binned KR balanced *cis* observed/expected matrix was extracted from “.hic” files. For each interaction bin-bin pair separated by a given genomic distance (*s*), we computed the product of two EV1 values corresponding to the two bins. We then calculated the Spearman correlation coefficient *R* between EV1 products and the observed/expected values of all bin-bin pairs that are separated by *s*. *R(s)* was then set to demarcate the level of compartmentalization for genomic distance *s*. To generate the *R(s)* curve across different genomic distances, we computed *R* when s equals 100kb, 200kb, 300kb……125Mb. *R(s)* curve of each chromosome were then averaged to generate the genome wide *R(s)* curve. *R(s)* curve of individual biological replicates and merged samples for both untreated control and auxin treated samples were computed independently across all cell cycle stages. For interactions close to the diagonal of contact maps, well-compartmentalized regions, i.e. interactions between bins from the same type of compartments (A-A or B-B) tend to display high observed/expected values and positive (>0) EV1 products, whereas interactions between bins from different types of compartments (A-B or B-A) tend to exhibit low observed/expected values and negative (<0) EV1 product. Thus, *R* tends to be high in well-compartmentalized regions. At weakly-compartmentalized regions, interactions between bins tend to be low also when distant from the diagonal regardless of whether the two bins are from the same type of compartment or not. Thus, *R* tends to be low in weakly-compartmentalized regions.

#### Interactions between local B-B or A-A compartments

To quantify the interactions between closely positioned compartments, we adopted a high resolution (50kb binned) A/B compartment profile from the mid-G1 untreated control sample as reference. To measure interactions between local B-B compartments (tier1, with one A-type compartment in between), we extracted genomic coordinates of all A-type compartments from the reference A/B compartment file. For each A-type compartment, we computed the average observed/expected values between the 250kb region up-stream of the start site and the 250kb region down-stream of the end sites across all cell cycle stages in both untreated control and auxin treated samples as well as G1-phase samples treated with triptolide. The resulting observed/expected values were denoted as interaction strengths between each closely spaced B-B compartment pair. We also computed more distally separated tier2 (with two A-type compartment in between) B-B interactions. For each given tier2 B-B compartment pair, we computed the average observed/expected values between the 250kb region up-stream of the start site of the first A-type compartment and the 250kb region down-stream of the end site of the second A-type compartment. A similar approach was taken to calculate tier3 (with three A-type compartment in between) and tier 4 (with four A-type compartment in between) B-B interaction. Interactions between tier1 to 4 A-A compartment pairs were calculated using the same approach.

For distance matched random controls, we randomly selected 500 A- or B-type compartments and shuffled their genomic coordinates for each entry using the “shuffle” function of bedtools^42^. Randomly shuffled bed files were used as input to compute the interaction strengths between 250kb up-stream and down-stream flanking regions for each entry.

#### Loop calling and post-processing

Chromatin loops were called using a previously described HICCUPS method with modifications^31^. The following steps were taken to generate unique non-redundant lists of loops in untreated controls and auxin treated samples across all cell cycle stages. (1) We used HICCUPS to call preliminary loops on the untreated control samples at each cell cycle stage using 10kb binned matrices with the Juicer_tool_1.13.02. The inner and outer diameters of the donut filter was set to be 4bins and 16bins respectively and an FDR of 0.2 was adopted. (2) We repeated the above step on 10kb binned auxin treated samples across all cell cycle stages with the exact same parameter set. (3) Loop calls from (1) and (2) were merged to generate a non-redundant loop list across all cell cycle stages and auxin treatment condition for 10kb matrices. (4) False positive calls introduced by exceptionally high outlier pixels were usually present in all samples irrespective of cell cycle stage and auxin treatment condition. To eliminate these artifacts, we removed pixels that were called in more than 6 of the 8 total samples. (5) In certain scenarios, pixels identified at different cell cycle stages or different auxin treatment conditions tended to cluster together. These clusters of pixels could actually be considered as one loop instead of many. Therefore, we implemented a method to merge these clustered pixels. To begin with, for a given loop in the non-redundant list from step (4), we recorded a value *q*_min_ which represents the lowest *q* value across all cell cycle stages in both untreated controls and auxin treated samples. We then ordered the loops in ascending order based on their *q*_min_. In this way, pixels at the top of the list were the most confident calls. We then focused on the top pixel and scanned through the rest of the list to identify pixels that were within a 20kb radius of the top pixel. If no additional pixel were found nearby, we then considered the top pixel as a loop by itself without pixel clustering. If pixels existed that fulfilled the above requirement, we then consider these pixels together with the top pixel as a loop cluster. Pixels within the loop cluster were removed from the non-redundant loop list. We then recalculated the centroid of the loop cluster and computed the distance *s* between the centroid and the far cluster edge. We then started a second round of pixel merging by scanning the rest of the list to identify pixels that are within a radius of 20kb + *s* from the centroid. Pixels within the loop cluster after the second collapsing step were removed from the non-redundant loop list. Next, we focused on the top pixel of the remaining list and repeated the above clustering steps, until no further pixels remained in the pixel list. After pixel merging, we generated a list of loop clusters which contained 1 or more pixels. For each loop cluster, we defined a cluster summit which was represented by the pixel with lowest *q*_min_. If a loop cluster only contained pixels called in the untreated controls but not in the auxin treated samples, we defined it as a “lost” loop. Conversely, if a loop cluster did not contain pixels from the control samples, it was defined as “gained” loop. The remaining loop clusters were categorized into “retained”, indicating that these loops were detected in both “-auxin” and “+auxin” samples. (6) We next performed step (1) through (5) on 25kb binned matrices with an inner donut filter diameter of 1 bin and outer donut filter diameter of 6 bins. FDR of 0.01 was adopted. (7) Loops called on 25kb binned matrices were then merged with those called on 10kb matrices. If a 25kb loop cluster overlaps with a 10kb loop cluster, the 25kb loop was dropped.

We noticed that some visually solid loop-like pixels were dropped by the HICCUPS due to lack of surrounding significant pixels. To recover these potentially false negative calls, we took advantage of our biological replicates. Specifically, we continued to complement our non-redundant loop calling list with the following steps. (8) For untreated control samples, we extracted all raw significant pixels from HICCUPS before clustering across cell cycle stages and combined them. (9) We then computed the donut FDR of the above pixels in all biological replicates across cell cycle stages in the untreated control samples through juicer_tools_1.13.02. (10) To determine if a pixel represented a loop, we implemented the below filters: <1>, For ana/telophase or early-G1 phase, we required that a pixel must display an FDR < 0.2 in both replicate-merged and individual biological replicates. <2>, For ana/telophase or early-G1 phase, a pixel was required to show an observed/donut-expected value of over 1.5 in replicate-merged and individual biological replicates. <3>, For ana/telophase or early-G1 phase, a pixel had to exhibit an observed value of >10 in replicate-merged and individual biological replicates. <4> For mid-G1 phase, the above three criteria had to be satisfied in at least 3 of the following 4: replicate-merged, biological replicate 1, 2 and 3. A pixel had to fulfill all the above filters to be viewed as valid in a given cell cycle stage, and it had to be valid in at least one post-mitotic cell cycle stage to be considered a valid loop for the untreated controls. (11) We then repeated step (8) through (10) to get a list of valid pixels in auxin treated samples. (12) We combined the valid pixels from untreated controls and auxin treated samples and filter out pixels with highest 5% observed/donut-expected values in prometaphase in untreated control samples as well as pixels with a distance of over 2Mb. (13) We further removed pixels that were overlapping or next to the loops identified in step (9). In this manner we obtained valid loops that had been previously missed by HICCUPS. (14) Finally, we performed step (5) on the remaining pixels at (13) to merge valid pixels that were clustered together. In total, we ended up with 16370 non-redundant loops across all samples.

#### Loop categorization based on CTCF/cohesin/CRE

We categorized loops into different classes based on whether ChIP-seq peaks of CTCF and cohesin and annotations of promoters or enhancers were present at their anchors. For a peak to intersect with a loop anchor, it had to have at least 1bp overlap with a 30kb region centered on the midpoint of the loop anchor summit. We employed the CTCF/cohesin co-occupied peak list and the peaks of H3K27ac (CRE) from our previous study ^1^. Our analysis focused on the following possibilities: (1) Two loop anchors harbor CTCF/cohesin co-occupied sites with neither harboring CREs. (2) Both loop anchors harbor CTCF/cohesin co-occupied sites with one anchor also harboring a CRE. (3) Both loop anchors each harbor a CTCF/cohesin co-occupied site and a CRE. (4) None of the loop anchors harbor CTCF/cohesin co-occupied sites but both contain CREs. (5) One loop anchor a harbors CTCF/cohesin co-occupied site and two anchors harbor CREs. Group (1) and (2) loops were defined as “structural loops”, loops from group (3) as “dual-function loops”, and loops from group (4) and (5) as “CRE loops”.

#### *K*-means clustering of loops

To measure the change of post-mitotic loop formation as well as the impact of CTCF depletion on loop formation, we defined a metric to measure the strength of each loop. For a given loop, we considered its summit pixel as well as 8 surrounding pixels and computed their observed/donut-expected values across cell cycle stages in both untreated controls and auxin treated samples. For a specific cell cycle stage and auxin treatment condition, the loop strength was recorded as the average of the observed/donut-expected values from the 9 pixels. To dissect the wildtype CRE loop reformation patterns after mitosis, we focused on the 3232 CRE loops that were detected in untreated control samples. We then computed the *z*-scores of loop strength across all cell cycle stages in untreated control samples and performed *k*-means clustering using the 3 post-mitotic time points. We were able to recover 4 loop clusters with distinct reformation kinetics. To assess the effect of CTCF depletion on cluster1 transient CRE loops, we attempted to further sub-categorize them using the loop strength from both untreated controls and auxin treated samples. We computed the *z*-scores of loop strength across all cell cycle stages in both untreated as well as auxin treated samples. We then performed *k*-means clustering using the 3 post-mitotic time points from both untreated controls and auxin treated samples.

#### Measuring the interplay between structural and CRE loops

To quantify the degree by which CRE loops are disrupted or supported by structural loops, we performed the following enrichment analysis. We first focused on structural loop interpolation. For a given cluster of CRE loops (e.g. cluster1-R), we defined two scenarios based on whether or not they are interpolated by structural loops. The two scenarios were: (1) neither of the two CRE anchors were covered by a structural loop (not interpolated) and (2) either one or both of the CRE anchors were covered by structural loops (interpolated). For each scenario, we constructed a 2 x 2 contingency table based on in which scenario a given cluster of CRE loops fell, and whether or not the rest of the CRE loops fell into that scenario. Odds ratios and *P*-values were computed with the Fisher’s exact test in R. Lastly, we defined for each cluster of CRE loops whether or not they were supported by structural loops.

A similar approach was taken to calculate the odds ratio and *P*-values for each CRE loop cluster at each scenario.

#### Domain calling

Domains were independently identified in untreated controls and auxin treated samples across all cell cycle stages, using the rGMAP algorithm (https://github.com/tanlabcode/rGMAP)^30^. To call domains in untreated control samples, we extracted 10kb binned KR balanced *cis* contact matrices from replicate-merged “.hic” files of each cell cycle stage, using the DUMP utility of juicer_tool (1.13.02)^31^. The contact matrices were used as input to feed in rGMAP for domain calling. For each cell cycle stage, we started by performing the following domain sweep: (1) Maximally 3 levels of domains were allowed (dom_order=3). (2) Maximal contact distances of 2Mb were allowed (maxDistInBin=200). This step generated a basic list of domains. To capture sub-domain like structures, we performed an additional domain sweep, which allowed a maximal contact distance of 500kb (maxDistInBin=50). Additional sub-domains were then added to the basal list to create a preliminary list of domains for each cell cycle stage. A similar approach was carried out to generate the preliminary domain list in the auxin treated samples. As a reference, we also performed rGMAP to call domains in late-G1 phase parental cell samples, using the same criteria as above.

#### Domain & boundary detection across cell cycle stages

For untreated control samples, domains called at each cell cycle stage were merged to create a total domain list. Domains from the late-G1 wildtype samples were added into the above list to serve as a reference. To ensure the validity of our domain calls, we established the following filters: (1) Domains called at prometaphase had to overlap with at least three domains identified in the subsequent four cell cycle stages (ana/telo, early-G1, mid-G1 and late-G1 from wildtype sample) to be considered valid. (2) Domains called at ana/telo, early-G1 or mid-G1 had to overlap with at least one domain identified in subsequent cell cycle stages to be considered valid. To claim that a domain detected in prometaphase is also present at a later cell cycle stage, we require that at least one domain exists in the later time point, whose upstream and downstream boundaries are within -/+ 8 bins of those of the original domain. We performed this step across all subsequent cell cycle stages to identify all potentially “overlapping” domains. If at least three subsequently identified domains overlap with our query domain, we then separately average the up- and down-stream boundaries of all “overlapping” domains to replace the boundaries of the original prometaphase domain. A similar approach was carried out in ana/telo, early-G1 and mid-G1 phase. These steps produced a list of high confidence domains that were detected across different cell cycle stages in the untreated control samples.

Next, we implemented a merging step to adjust boundary locations so that domains across cell cycle stages with highly similar boundaries would share a single consistent boundary (It’s noteworthy that possibilities still remain that two highly similar boundaries represent true biological differences instead of technical differences.). As a start, we generated an overall non-redundant boundary list from all domains. Boundaries were then sorted based on their genomic coordinates from 5’ to 3’. Starting from the first boundary, we swept throughout the rest of the boundaries on the same chromosome and removed boundaries that are less than 80kb away from the first boundary. We then, merged these boundaries into one and applied the mean of their genomic coordinates as the genomic coordinate of the final merged boundary. These boundaries were then removed from the overall boundary list. We then performed this step iteratively on the remaining boundaries until all boundaries were processed. The final averaged boundary coordinates were then reassigned to corresponding domains. For a boundary shared by multiple domains, the time point of emergence of this boundary is determined by the earliest associated domain.

The same approach was carried out to process domains and boundaries identified in the auxin treated samples across all cell cycle stages.

#### Insulation score profiling

Insulation scores were computed as previously described^43^. Briefly, we implemented a 12bin x 12bin window, which slides along the diagonals of the 10kb binned KR balanced contact matrices. The sliding window was set to be one bin away from the diagonal. Genomic regions with low read counts (<12 counts) were discarded from the analysis. Windows interrupted by the starts or ends of chromosomes were also discarded. For each 10kb bin, the sum of read counts of each window was then normalized to the chromosomal average and log_2_ transformed. A pseudo read count was added to the chromosomal mean as well as each window before log transformation.

We noticed that in a few cases domain boundaries were shifted by several bins from the local minima of insulation, and thus did not accurately reflect the “real” boundary position. To solve this issue, we fine-tuned boundary positions such that boundaries were adjusted to the local minima of insulation. This adjustment was performed on untreated control samples. For each given boundary, we defined a wiggle room by sectioning a -6bin to +6bin genomic region that centered around the boundary. We then recorded the mid-G1 phase insulation scores of each bin within the wiggle room. The bin with lowest insulation score for mid-G1 phase was defined as the final position of the boundary, representing local minima of insulation scores. The adjusted boundary locations were then re-assigned to their corresponding domains. After boundary adjustment, domains smaller than 100kb were filtered out to eliminate spurious domains. In some outlier cases, boundaries after adjustment ended up being extremely close to each other (within 20kb). We therefore implemented a final step to merge boundaries using the same approach as described above.

#### Integration of domains and boundaries from untreated control and auxin treated samples

After boundary position adjustment, we obtained an intermediate list of domains and boundaries that were detected at each cell cycle stage for the untreated control samples. We then added domains and boundaries from “+auxin” samples to this list to create a final complete domain list before quality check. We carried out the flowing merging steps: For a given domain in the auxin treated samples, if both of its boundaries were less than 80kb away from the up- and downstream boundaries of a “-auxin” domain, we then considered these two domains as “overlapping” and recorded the boundary coordinates of the “-auxin” domain in the final list. If the upstream (but NOT downstream) boundary of the “+auxin” domain was less than 80kb away from any boundaries in the “-auxin” list, we then considered this “+auxin” domain a new domain and recorded the “-auxin” boundary coordinate as the upstream boundary for this “+auxin” domain. Similarly, if the downstream (but NOT upstream) boundary of the “+auxin” domain was less than 80kb away from any boundaries in the “-auxin” list, we considered this “+auxin” domain as a new domain and recorded the “-auxin” boundary coordinate as the downstream boundary for this “+auxin” domain. Finally, if both upstream and downstream boundaries of the “+auxin” domain were more than 80kb away from boundaries in the “-auxin” list, we considered this “+auxin” domain as a new domain and recorded its own boundary coordinates in the final list of domains.

#### Domain quality check and aggregated domain analysis (ADA)

We noticed that in some rare cases, domains spanning large low-mappable regions were also called by the algorithm. To filter out low confidence domains, we implemented an aggregated domain analysis which measures the ratio between interactions just inside the domain and interactions just outside the domain. We computed ADA scores on the 10kb binned KR balanced observed/expected contact matrices as previously reported with modifications^1, 12^. For each domain, the start and end coordinates were recorded as i x 10000 and j x 10000 respectively. Therefore, we could use (i, j) to mark the position of the corner pixel. We then marked our 4 horizontal stripes and 4 vertical stripes that were just inside the domains. The positions of the horizontal inner stripes are: [i+1, j-8: j-4], [i+2, j-7:j-3], [i+3, j-6:j-2] and [i+4, j-5:j-1] respectively. The positions of vertical inner stripes are: [i+1:i+5, j-4], [i+2:i+6, j-3], [i+3:i+7, j-2] and [i+4:i+8, j-1] respectively. We then also marked out additional 4 horizontal and 4 vertical stripes that were outside of the domain. The positions of horizontal outer stripes were [i-8, j-17:j-13], [i-7, j-16:j-12], [i-6, j-15:j-11] and [i-5, j-14:j-13], respectively. The positions of vertical outer stripes were [i+10:i+14, j+5], [i+11:i+15, j+6], [i+12:i+16, j+7] and [i+13:i+17, j+8], respectively. The inner stripes and outer stripes had the same genomic separations. We computed the sum of observed/expected values for pixels within inner stripes and then divided this value by the sum of observed/expected values for pixels within outer stripes. The final result was log_2_ transformed and recorded as the ADA score of the domain. Note that domains smaller than 150kb was filtered to minimize the possibility of outer stripes stretching into another domain.

To eliminate domains covering low-mappable regions, we placed the following filters: (1) For a given domain, we examined all pixels in the inner and outer stripe regions. If any pixel displayed an observed/expected value of over 30, we dropped this domain from further analysis. This step was to filter out high outlier pixels that are usually associated with low-mappable regions. (2) For a given domain, if either or both of the two outer stripe regions contained less than 5 non-zero pixels, the domain was dropped from further analysis. (3) For a given domain, if either or both of the two inner stripe regions contain less than 10 non-zero pixels, the domain was dropped from further analysis. To further ensure the validity of our domain calls, we also implemented a dynamic filter. This filter was established based on the rationale that true domains would gradually become stronger after mitotic exit and thus their ADA scores would be higher in post-mitotic time points compared to prometaphase. Specifically, we require that for a domain to be valid, at least 1 of the 6 post-mitotic samples (“ana/telo -auxin”, “ana/telo +auxin”, “early-G1 -auxin”, “early-G1 +auxin”, “mid-G1 -auxin”, “mid-G1 +auxin”) had to show at least a 1.25-fold ADA score enrichment compared to both prometaphase samples (“prometa -auxin”, “prometa +auxin”).

#### Dynamic clustering of boundaries

We quantified boundary strength as follows: We selected a - 120kb to +120kb region centered on a boundary of interest and searched for the highest insulation score within this 240kb region. This maxi-IS value was then unlogged and subtracted by the insulation score (unlogged) at the boundary itself. The resulting ΔIS was then denoted as the strength of the target boundary.

To examine the reformation dynamics of boundaries and measure the effect of CTCF depletion on the dynamic boundary formation after mitosis, we performed *k*-means clustering on the ΔIS of all boundaries across all 8 samples (4 cell cycle stages in both untreated control and auxin treated samples). Specifically, for each boundary, we computed the *z*-scores of their ΔIS across all 8 samples. We then performed k-means clustering on the *z*-scores of 6 post-mitotic samples (ana/telo, early-G1 and mid-G1 x 2 treatment conditions). We found that when we chose *k*=5 clusters, we were able to recover the most biologically interpretable clusters. Note, cluster5 was mostly spurious boundaries and thus were excluded from the analysis.

#### PCA based interrogation of chromatin state transition at boundaries

To assess histone modification features associated with different clusters of boundaries, we adopted a PCA based approach as previously described ^44^. For each target boundary, we selected a −50kb to +50kb genomic region and sectioned it into 10bins (10kb/bin). To assess chromatin state transition, we adopted the two histone marks H3K36me3 and H3K27me3, the former and latter representing transcriptionally active and inactive chromatin, respectively. We then calculated the mean G1E-ER4 ChIP-seq signals of these two marks (from asynchronously growing cells) in each 10kb bin, using the UCSC toolkit (BigWigAverageOverBed). The ChIP-seq intensities of these marks were organized into two matrices such that each row represents the 100kb region around a boundary, and each of the 10 columns represents a 10kb bin. The columns were ordered based on their genomic positions from upstream to downstream. Each column was then normalized to the column sum such that the ChIP-seq intensity values from each column add up to 1. After normalization, the two matrices of H3k27me3 and H3K36me3 were stitched horizontally yielding a final matrix with 20 columns. We then applied principal component analysis (PCA) on the final matrix using the R package (“prcomp”). We noticed that PC1 was able to accurately describe the transition of chromatin states in a way that boundaries with either highest or lowest PC1 projection values were typically at chromatin transition points (5’ active → 3’ inactive or 5’ inactive → 3’ active), whereas boundaries with median level PC1 projection values were not.

#### PolII ChIP-seq processing and peak calling

Reads were aligned against the mm9 reference genome using Bowtie2 (v2.2.9) with default parameters and soft clipping allowed (“--local”)^45^. Alignments with MAPQ score lower than 10 and PCR duplicates were removed using SAMtools (v0.1.19)^46^. Reads aligned to mitochondria, random contigs and ENCODE blacklisted regions were also removed for downstream analysis using BEDtools (v2.27.1)^42^. Peaks were called using MACS2 (v2.1.0) with default parameters and a 0.01 q-value cutoff. Fragment pileup and local lambda track files in bedGraph format were created during MACS2 peak calling and normalized to one million reads per library (“callpeak --bdg --SPMR”)^47^. The latter was track was subtracted from the former using MACS2 (“bdgcmp -m subtract”), negative values were reassigned as zeros, and converted to bigwig format for visualization using the UCSC Toolkit (“bedGraphToBigWig”). Finally, a non-overlapping union set of peaks was created by merging peaks in all replicates using BEDtools such that all peaks that overlap by at least 1bp were merged.

#### Identification of active genes

Active genes were called based on the overall PolII ChIP-seq peak list with the following filters: (1) The TSS of a gene had to overlap with at least 1 positive PolII ChIP-seq peak. (2) The length of a gene had to be over 1kb to ensure that enough reads were obtained over the gene body (+500bp from TSS to TES) and discernible from the reads at the TSS. (3) We further filtered out the genes with the lowest (10%) H3K27ac signal or ATAC signal at the promoter regions (−250bp to 250bp of TSS, data from asynchronous G1E-ER4 cells)^48, 49^. A PolII ChIP-seq peak at the TSS does not necessarily mean that the corresponding gene is active. In certain cases, inactive genes positioned closely downstream of the 3’ UTR of active genes could also display positive PolII signals at their TSS, potentially leading to false assignment as active gene. Therefore, we filtered out genes with low H3K27ac or ATAC signal to ensure that the genes were within “open” chromatin and more likely to be active. (4) The PolII ChIP-seq signal (+500bp from TSS to TES) of at least 1 of the six post-mitotic samples (“early-G1 -auxin”, “early-G1 +auxin”, “mid-G1 -auxin”, “mid-G1 +auxin”, “late-G1 -auxin”, “late-G1 +auxin”) had to be ≥ 1.5 fold that of the two prometaphase (“-auxin” and “+auxin”) samples.

#### PCA based interrogation of post-mitotic gene activation pattern

PCA was performed separately on PolII ChIP-seq signals from control and auxin treated samples. For untreated samples, we computed the replicate-merged PolII ChIP-seq signals (+500bp from TSS to TES) of each active gene across all cell cycle stages, using the UCSC toolkit (BigWigAverageOverBed). The PolII signals from each cell cycle stage were then normalized such that they sum up to 1. PCA was performed on the last 3 cell cycle stages using the R package (prcomp). As described above, the PC1 values of each gene describe the “spikiness” of their post-mitotic reactivation pattern. We set the direction of PC1 projection values such that genes with high (positive) PC1 values were the most “spiky” after mitosis, whereas genes with low (negative) PC1 values displayed a gradual increase of PolII ChIP-seq signal after mitosis. The same procedure was performed on auxin treated samples. The PC1 values of each gene from control and auxin treated samples were highly correlated, suggesting that the post-mitotic transcriptional spiking was maintained after CTCF depletion.

#### Differential gene expression analysis

Gene expression levels were assessed by the number of PolII ChIP-seq read counts over the gene body (+500 from TSS to TES). To measure differential gene expression after CTCF depletion during mitotic exit, we first extracted raw PolII read counts over gene bodies from the bam files of each individual biological replicate using the “multicov” function of bedtools. This step was performed on the 3 post-mitotic cell cycle stages in both “-auxin” and “+auxin” samples. DESeq2 was adopted to perform differential expression analysis between “-auxin” and “+auxin” samples for each post-mitotic cell cycle stage independently. Raw read PolII ChIP-seq read counts were used as input for DESeq2 with default parameters^50^. *P.adj* cutoff of 0.05 and fold change cutoff of 1.25 were adopted to call differentially expressed genes for each post-mitotic cell cycle stage. A gene was considered differentially expressed if it was significantly different in at least one post-mitotic time point. In total, we identified 426 differentially expressed genes during mitotic exit after CTCF depletion. To determine whether these genes were up- or down-regulated over time, we performed *k*-means clustering using the log_2_FC output of DESeq2 across early-, mid- and late-G1 phase. Finally, we identified 223 genes which were down-regulated after CTCF depletion, and 203 genes which were up-regulated after CTCF depletion during the mitosis to G1-phase transition.

#### ABC model to predict functional enhancers of active genes

To predict enhancers of active genes and establish E-P and P-P connections, we adopted a recently proposed ABC model (https://github.com/broadinstitute/ABC-Enhancer-Gene-Prediction) ^32^. To simulate enhancer activity, we used H3K27ac ChIP-seq and ATAC-seq signals from asynchronously growing G1E-ER4 cells. These datasets were used in combination with 6 replicate-merged Hi-C datasets in this study (ana/telophase “-auxin”, ana/telophase “+auxin”, early-G1 “-auxin”, early-G1 “+auxin”, mid-G1 “-auxin”, mid-G1 “+auxin”) to predict enhancers in each of the 3 post-mitotic cell cycle stage with or without auxin treatment independently. We called E-P and P-P connections in each sample when the ABC score threshold equals to 0.01, 0.02, 0.03, 0.04 and 0.05 respectively. Higher the ABC score thresholds resulted in fewer but higher confidence connections. Note that we identified on average ∼1.86 (fewer than the recommended 3) enhancers per gene when the ABC score threshold was set to 0.04, suggesting that 0.04 is a relatively stringent threshold. For a given ABC threshold (e.g. 0.01), we combined the predicted connections in each sample to generate an overall non-redundant list of high confidence E-P and P-P pairs. Each E-P or P-P pair was then assigned to genes with different responses (up-reg, down-reg and non-reg) to CTCF depletion.

#### Differential interaction analysis between E-P contacts

Differential interaction analysis was carried out on E-P contacts called by ABC modeling with ABC score threshold set to 0.04. We adapted the E-P interaction strength values from ABC modeling and used as the input for LIMMA R package. Since the trend of CRE contact changing was overwhelmingly consistent between early and mid-G1 phase samples, we treated the samples from these two time points as equal biological replicates. Thus, we had 5 biological replicates (2 from early-G1 and 3 from mid-G1) for control and auxin treated samples. LIMMA was used to determine differentially interacting E-P contacts. *P* values were calculated with the eBayes function within LIMMA and adjusted with the Benjamini–Hochberg method. An FDR of 0.1 was used to call significantly strengthened or reduced E-P contacts.

#### Aggregated plots for loops, domains and compartments

Aggregated plots were generated using the python package “Coolpup”^51^. For unscaled aggregated peak analysis (APA), loops smaller than 100kb were removed from the plots to avoid influence from pixels close to the diagonal. For unscaled aggregated plots of compartment transition points, compartments smaller than 300kb were removed from the plots, again to minimize influence from pixels near the diagonal.

#### Data availability

HiC and PolII ChIP-seq data are deposited into GEO data base with accession number GSE168251. Additional external ChIP–seq data of histone modifications on asynchronous cells are available at: H3K27ac (GSE61349) ^48^, H3K4me1 (GSM946535) ^52^, H3K4me3 (GSM946533) ^52^, H3K36me3 (GSM946529) ^52^, H3K27me3 (GSM946531) ^52^, H3K9me3 (GSM946542) ^52^. Additional external data of CTCF and Rad21 before and after CTCF depletion in asynchronous G1E-ER4 cells are available at GSE150418 ^53^.

---

**Extended Data Figure 1.**
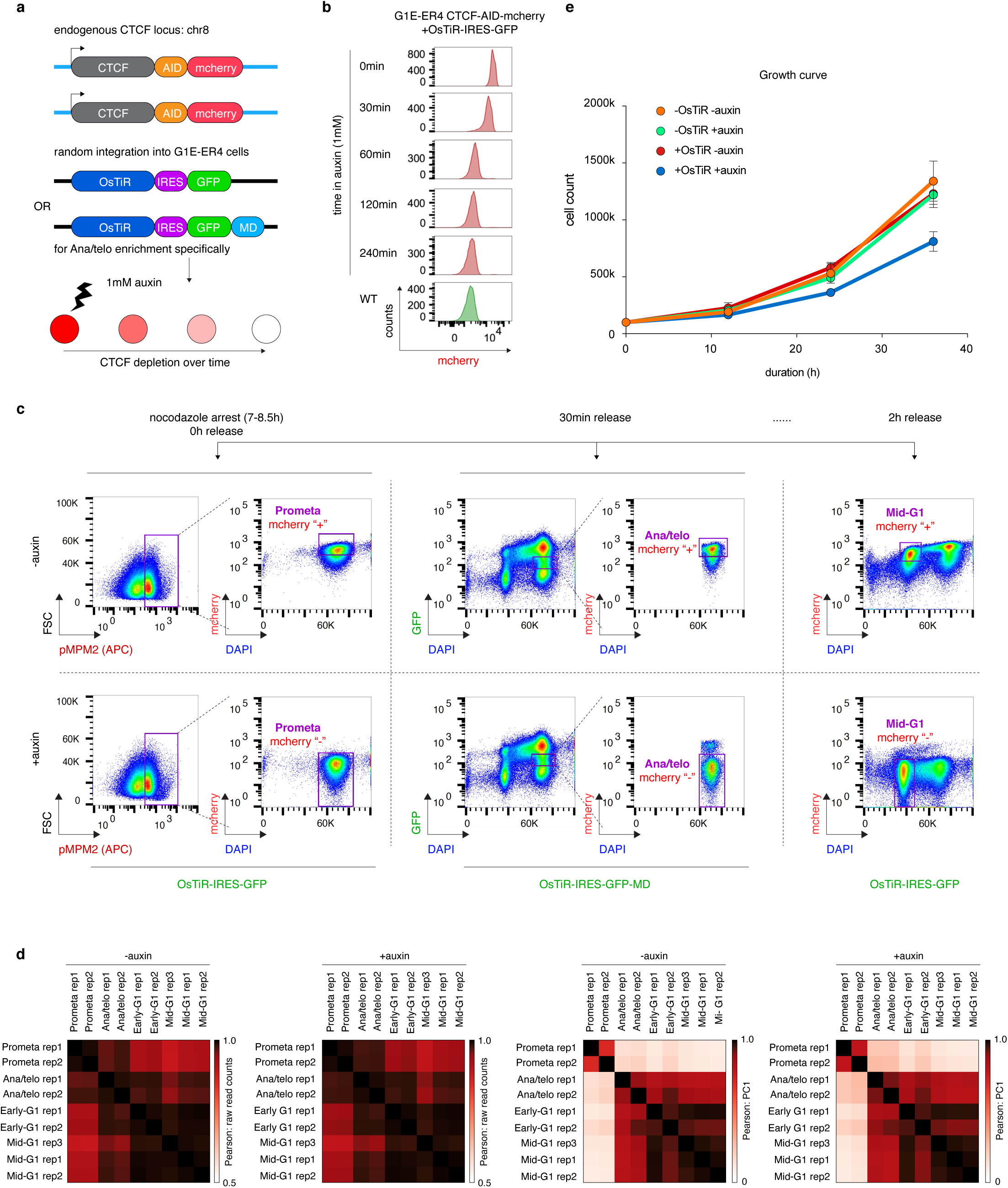
Engineering, purification, and characterization of cell lines. **a**, Schematic showing the construction of G1E-ER4 CTCF-AID-mCherry cell line and ectopic expression of Os-TIR-IRES-GFP. Os-TIR-IRES-GFP was used for prometaphase, early- and mid-G1 phase. To enable enrichment of ana/telophase cells, we specifically over expressed Os-TiR-IRES-GFP-MD. **b**, Flow cytometry plot showing the acute depletion of mCherry signal in asynchronous cells upon auxin treatment. Flow plots are representative of two independent experiments. **c**, FACS plots and gates (purple boxes) used for purification of mitotic and post-mitotic populations with and without CTCF. One set of plots representative of two independent biological replicates is show. **d**, Heatmaps showing Pearson correlations among Hi-C samples based on 100kb binned raw read counts and eigenvector 1 values, respectively. Note that samples with or without auxin treatment were separately plotted. **e**, Growth curves showing cell proliferation with or without auxin treatment. Note, cells devoid of OsTiR were included as controls. Error bar denote SEM (n=3).

**Extended Data Figure 2.**
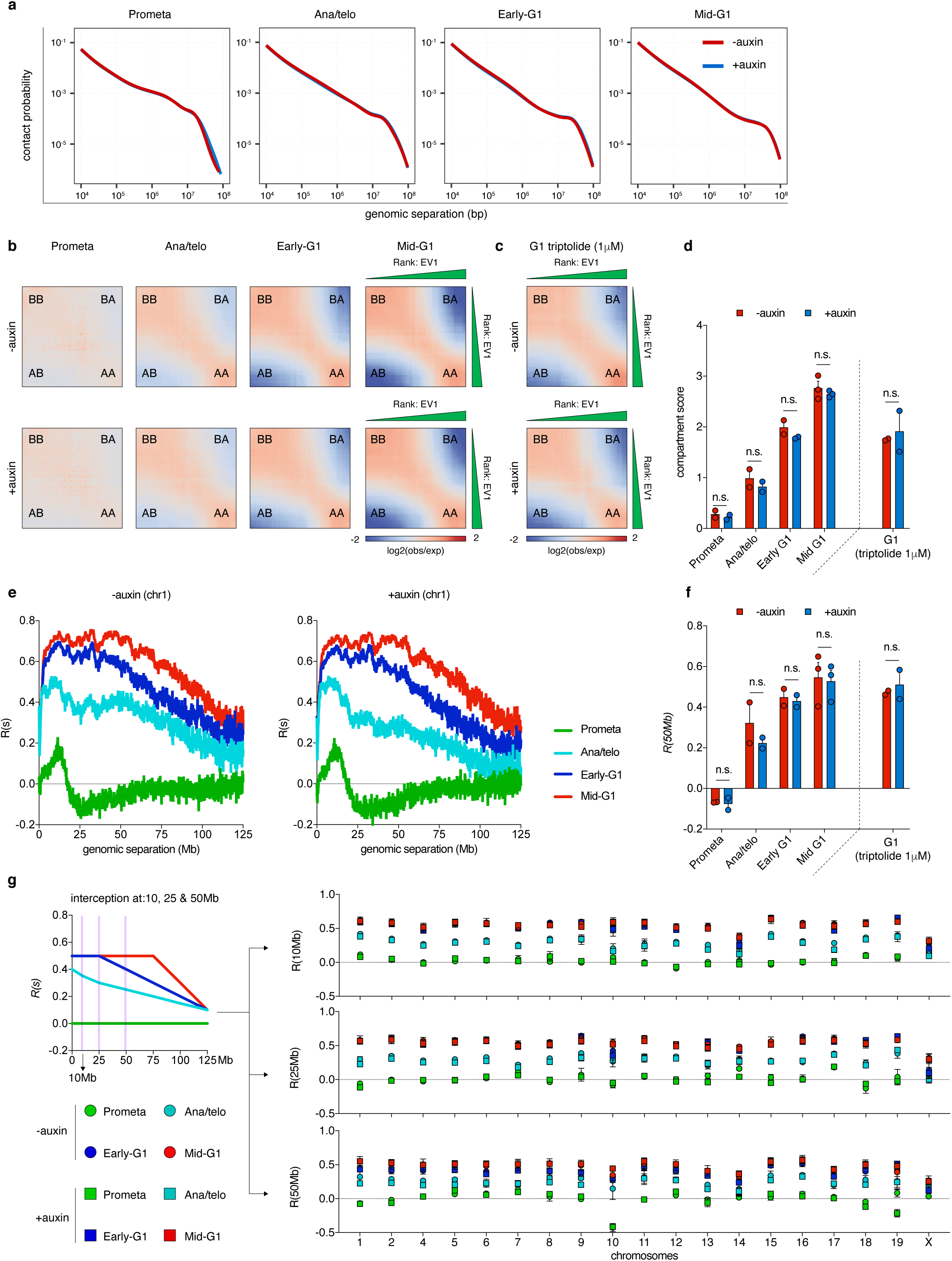
Global compartment re-establishment is unperturbed by CTCF depletion after mitosis. **a**, Chromosome averaged distance dependent contact frequency decay curves across cell cycle stages in control and auxin treated samples. **b**, Saddle plots showing compartment strengths across cell cycle stages in untreated and auxin treated samples. **c**, Saddle plots showing compartment strength in G1 phase cells after triptolide treatment. **d**, Bar graphs showing compartment scores for each cell cycle stage in untreated and auxin treated samples. Compartment scores of triptolide treated G1 samples were plotted on the right. Error bars denote SEM. Two-sided student’s t test was applied. **e**, Line graphs showing the level of compartmentalization *R* vs. genomic separations *s* for chromosome 1 for each cell cycle stage in both untreated and auxin treated samples. The gradual flattening of curves as cells progress towards G1 suggests expansion of the plaid-like compartmental interaction patterns from diagonal proximal regions to diagonal distal regions. **f**, Bar graphs showing the level of compartmentalization at 50Mb *R(50Mb)* for each cell cycle stage with or without CTCF. *R(50Mb)* of the triptolide treated G1 samples were plotted on the right. Error bars denote SEM. Statistical test: two-sided student’s t test. **g**, Left: Cartoon line plot of *R(s)* across different time points showing a series of intersections at 10Mb, 25Mb and 50Mb. Right: Replicate averaged *R(s)* of each individual chromosome across all cell cycle stages in both untreated and auxin treated samples when *s* equals to 10Mb, 25Mb and 50Mb respectively.

**Extended Data Figure 3.**
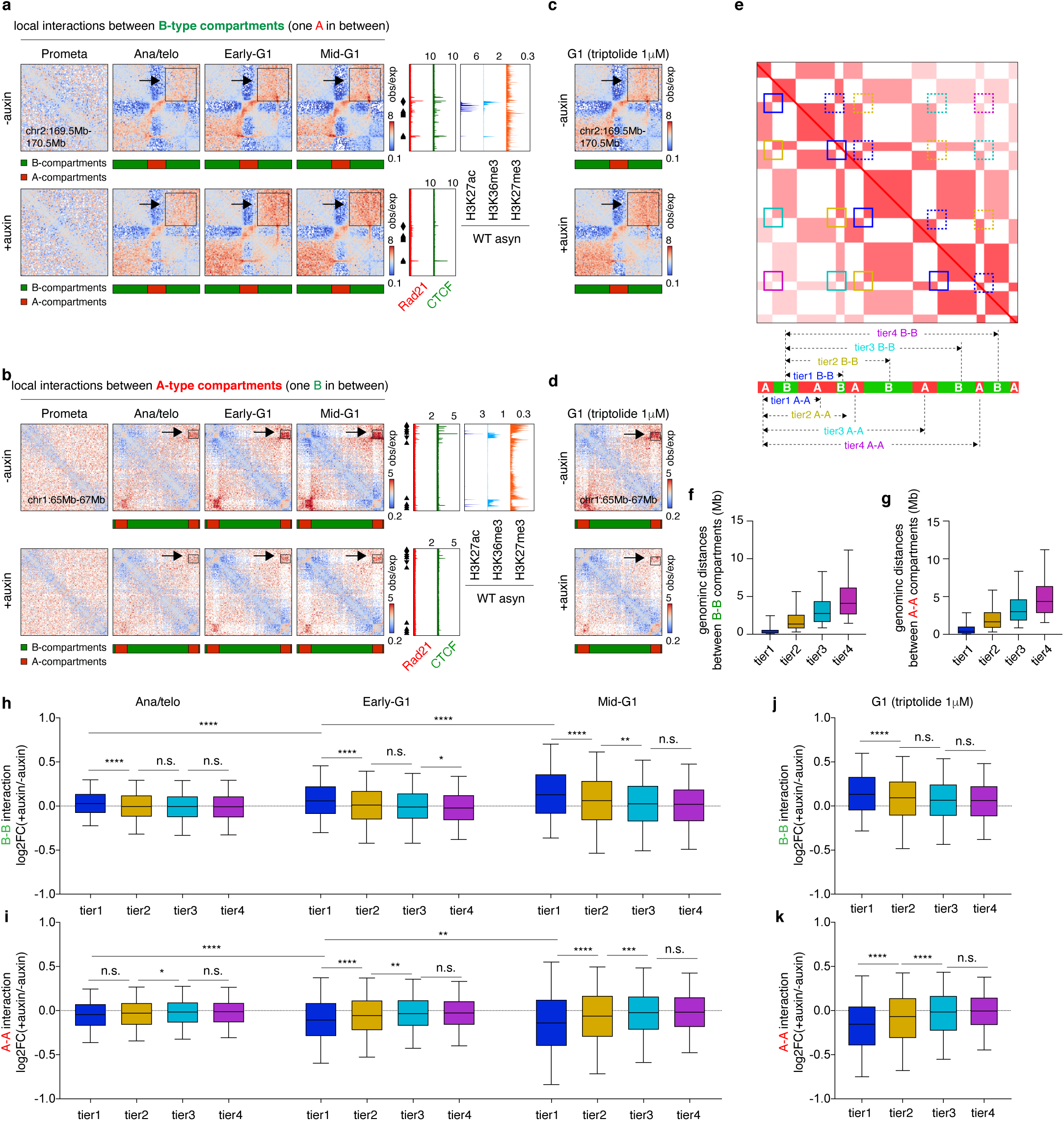
CTCF depletion alters local chromatin compartmentalization. **a**, Additional example of enhanced local B-B interactions after CTCF depletion. Bin size: 10kb. Arrows and boxes highlight the increased local B-B interactions after CTCF depletion across post-mitotic cell cycle stages. Tracks of CTCF and Rad21 with or without auxin treatment as well as H3K27ac, H3K36me3 and H27me3 are from asynchronously growing G1E-ER4 cells. **b**, Additional example of reduced local A-A interactions after CTCF depletion. **c**, KR balanced Hi-C contact matrices showing the same region as in (**a**) in G1 phase cells after triptolide treatment. **d**, KR balanced Hi-C contact matrices showing the same region as (**b**) in G1 phase cells after triptolide treatment. **e**, Schematic of the checkerboard pattern of compartments. Tier1-4 B-B interactions denote contacts between B-type compartments interspersed with 1-4 A-type compartments and are demarcated by dotted boxes. Tier1-4 A-A interactions denote contacts between A-type compartments interspersed with 1-4 B-type compartments and are demarcated by solidly lined boxes. **f and g**, Boxplots showing the distances between B-B or A-A interactions, respectively from different tiers. For all boxplots, central lines denote medians; box limits denote 25th–75th percentile; whiskers denote 5th–95th percentile. **h**, Boxplots showing the log_2_ fold change of B-B interactions from different tiers, upon CTCF loss. Similar comparisons are shown across all post-mitotic cell cycle stages. Comparisons between tier1 B-B interactions across cell cycle stages suggest the progressively amplified CTCF depletion induced gains of B-B interactions after mitosis. For all boxplots, central lines denote medians; box limits denote 25th–75th percentile; whiskers denote 5th–95th percentile. * *P* < 0.05, ** *P* < 0.01, *** *P* <0.001 and **** *P* <0.0001. Statistical comparisons within single time points are based on two-sided Mann-Whitney U tests. Comparisons across time points were computed by Two-sided paired Wilcoxon signed-rank test. **i**, Similar to (**h)**, displayed are log_2_ fold changes of A-A interactions from different tiers, upon CTCF loss. **j**, Boxplots showing the log_2_ fold change of B-B interactions from different tiers upon CTCF loss after triptolide treatment. For all boxplots, central lines denote medians; box limits denote 25th–75th percentile; whiskers denote 5th–95th percentile. * *P* < 0.05, ** *P* < 0.01, *** *P* <0.001 and **** *P* <0.0001. Two-sided Mann-Whitney U test. **k**, Similar to (**j**), shown are log_2_ fold changes of A-A interactions from different tiers upon CTCF loss after triptolide treatment.

**Extended Data Figure 4.**
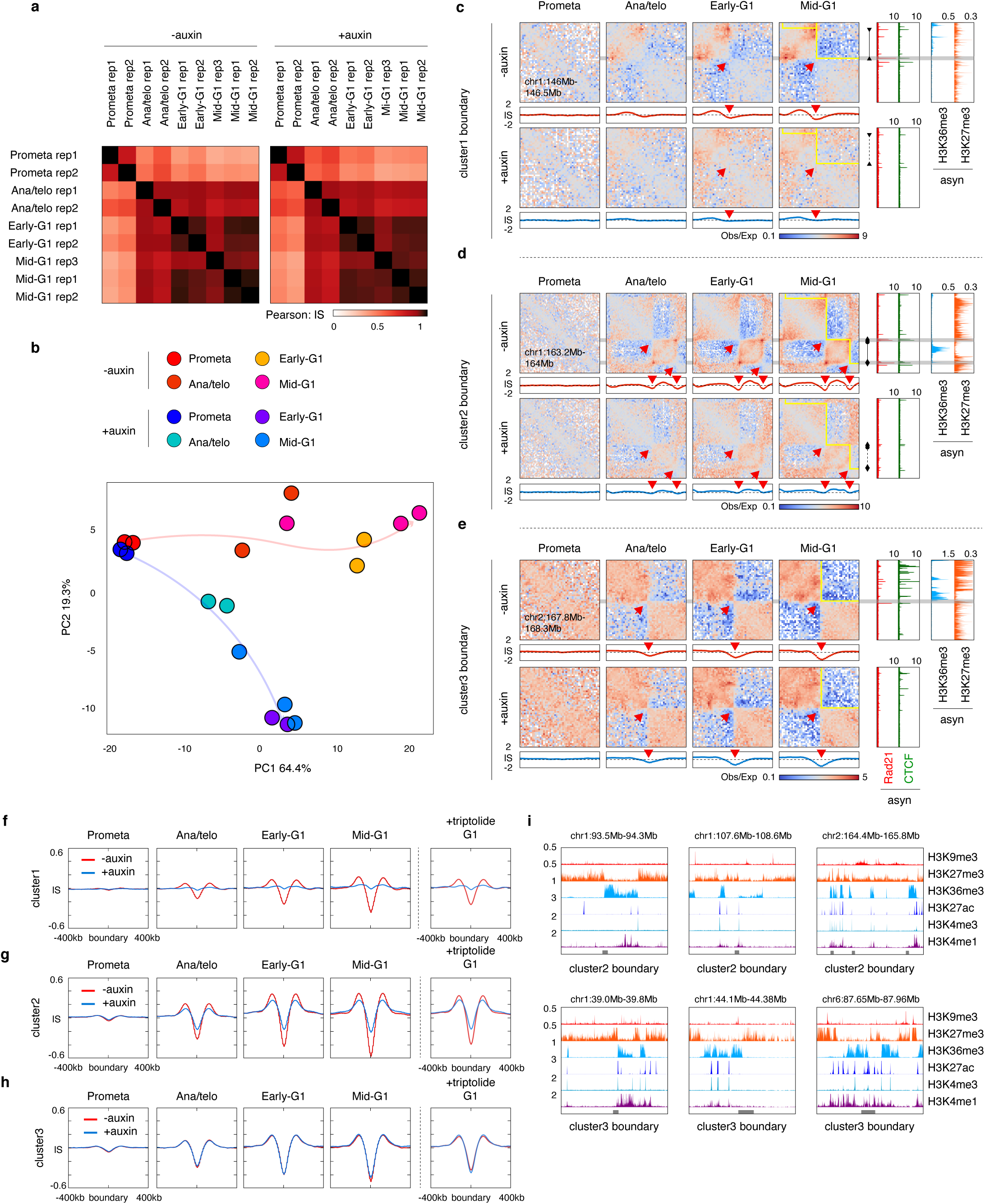
Characterization of boundary re-establishment after mitosis upon CTCF loss. **a**, Pearson correlations of boundary insulation scores between biological replicates in untreated or auxin treated samples. **b**, PCA analysis of the post-mitotic boundary reformation trajectories of with or without CTCF depletion. **c**, KR balanced Hi-C contact matrices and corresponding insulation score tracks of a representative region containing a cluster1 boundary across cell cycle stages in control and auxin treated samples. Bin size: 10kb. Read arrows indicate domain boundaries. Yellow lines denote TADs identified by rGMAP. Tracks of CTCF and Rad21 with or without auxin treatment as well as H3K36me3 and H27me3 are from asynchronously growing G1E-ER4 cells. Black arrow heads denote CTCF motif orientation, and dotted line demarcates loss of loop upon auxin treatment. **d & e**, Similar to (**c**), KR balanced Hi-C contact matrices showing representative regions containing cluster2 and 3 boundaries respectively. **f-h**, Left panel: Meta-region plots of insulation score profiles centered on cluster1, 2 or 3 boundaries, across cell cycle stages in control and auxin treated samples. Right panel: Meta-region plots of insulation score profiles similar to the left in G1 samples after triptolide treatment. **i**, genome browser tracks showing the local histone modification profiles of representative cluster2 and 3 boundaries.

**Extended Data Figure 5.**
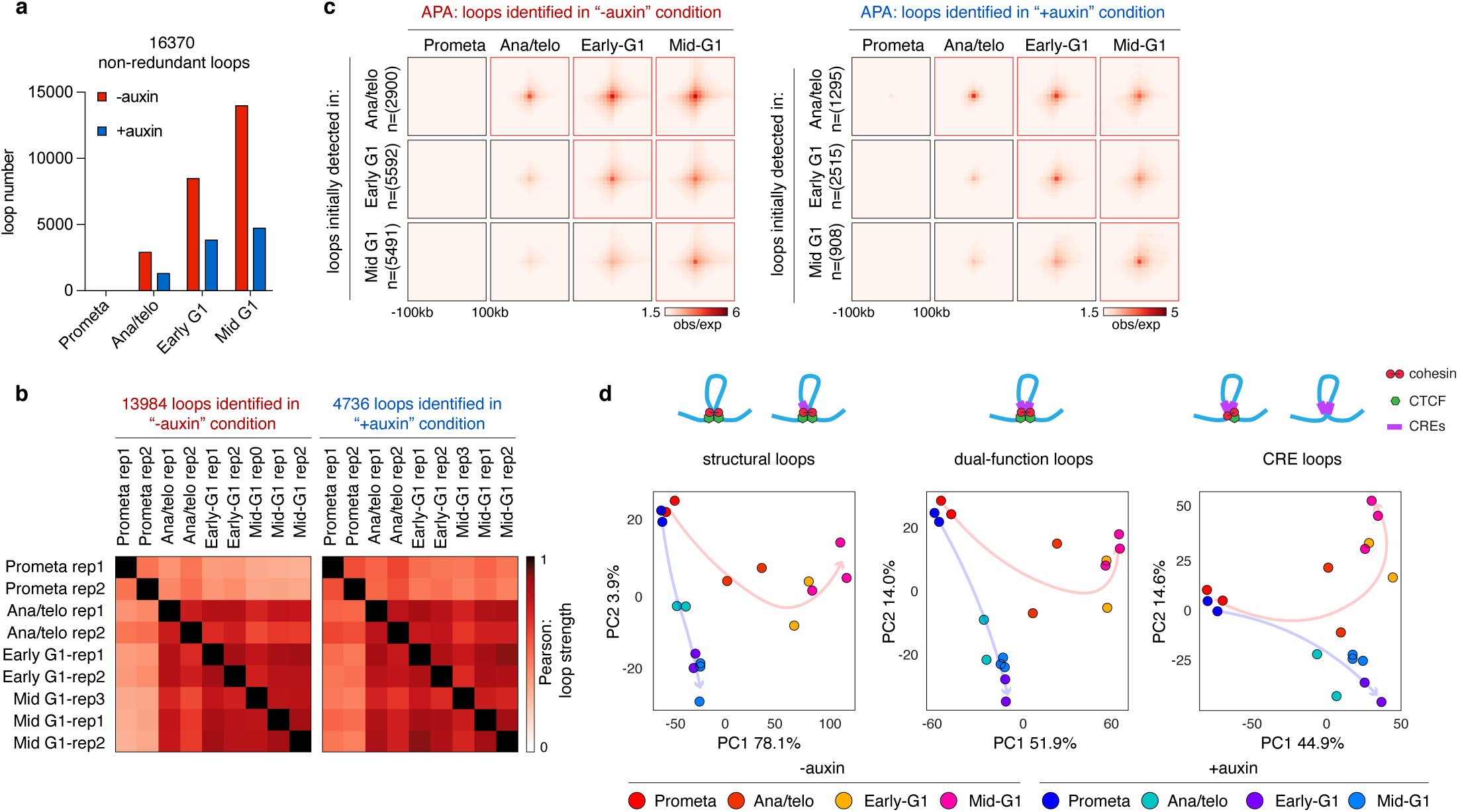
Distinct responses to CTCF depletion in chromatin loop reformation after mitosis. **a**, Bar graphs showing the number of loops called at each cell cycle stage in both untreated samples and auxin treated samples. **b**, Heatmaps showing the Pearson correlation of loops called in untreated and auxin treated samples, respectively. Pearson correlations were computed based on loop strength (obs/exp). **c**, Aggregated peak analysis (APA) plots showing time dependent emergence of loops called at each cell cycle stage in untreated and auxin treated samples. Bin size: 10kb. Notably, weak APA signals were observable before the loops were called, likely due to thresholding by the loop calling algorithm. **d**, PCA analysis showing the reformation trajectories of structural loops (left), dual-function loops (middle) and CRE loops (right) respectively, in untreated as well as auxin treated samples.

**Extended Data Figure 6.**
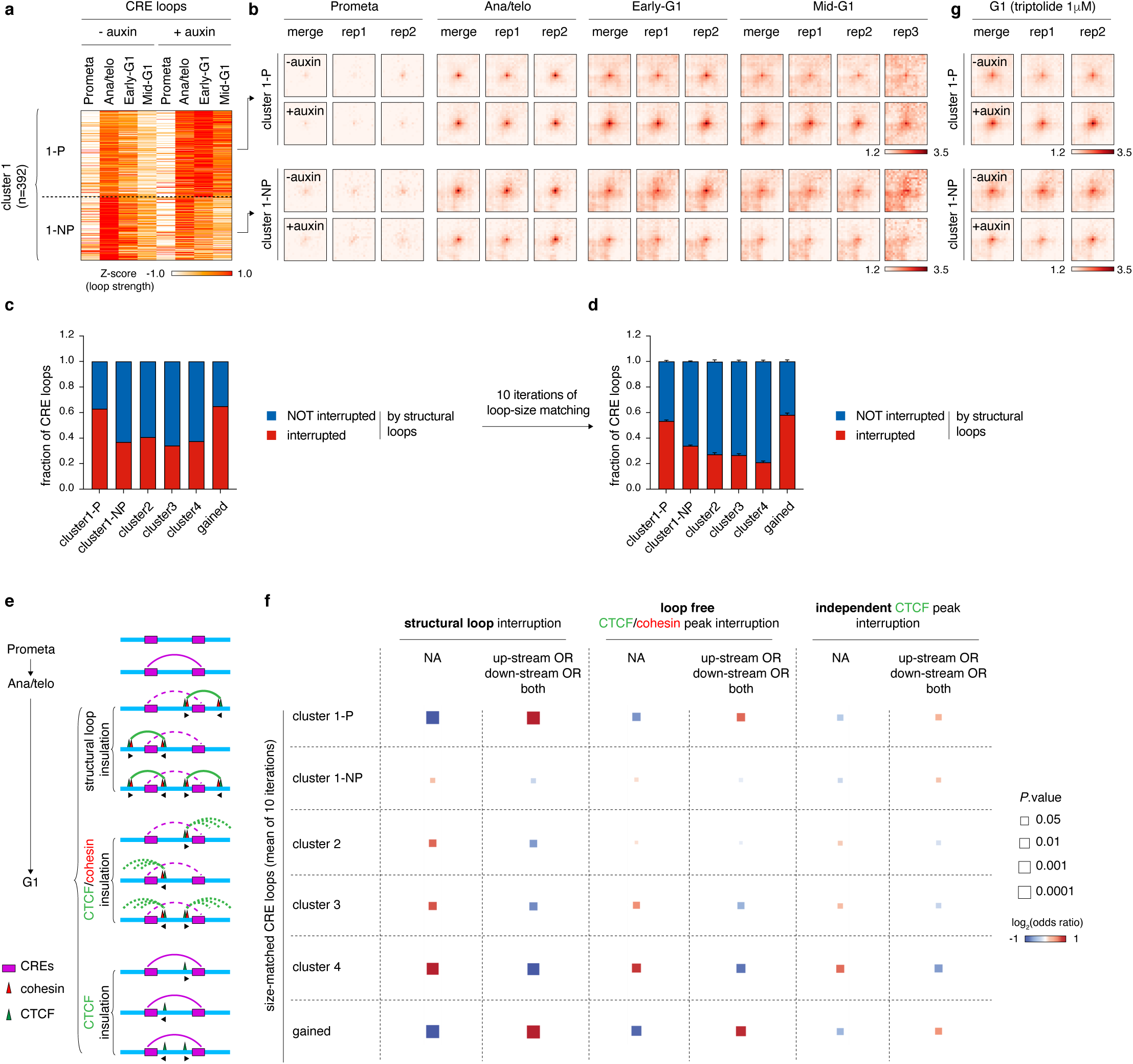
Transient CRE loops are terminated with the emergence of interfering structural loops. **a**, Heatmaps showing the sub-clustering of cluster1-P (persistent upon CTCF depletion) and cluster 1-NP (non-persistent upon CTCF depletion). **b**, APA plots of loop clusters from (**a**). Bin size: 10kb. Plots for the replicate-merged as well as each individual replicates are shown. **c**, Bar graphs showing the fractions of CRE loops from indicated clusters that were interrupted by structural loops. **d**, Bar graphs depicting the fraction of size-matched CRE loops from indicated clusters that were interrupted by structural loops. Error bars denote the SEM of 10 iterations of random size-matching operations. **e**, Schematic showing potential mechanisms (structural loops, CTCF/cohesin loop extrusion, and loop-independent CTCF) through which CTCF may exert its insulation function to disrupt transient post-mitotic CRE loops. **f**, Enrichment analysis of each sub-cluster of CRE loops controlled by mechanisms in (**e**). Left: relative enrichment analysis of whether CRE loops from each sub-cluster were interrupted by structural loops. Colors of the squares indicate log_2_ transformed odds ratio (Fisher’s exact test). Sizes of squares indicate the significance of enrichment (*p* value from Fisher’s exact test). Middle: similar to the left, showing relative enrichment of CRE loops from each sub-cluster to be interrupted by loop-free CTCF/cohesin co-occupied sites. Right: relative enrichment of CRE loops from each sub-cluster interrupted by cohesin and structural loop-independent CTCF peaks. **g**, APA plots showing the same pile-up Hi-C matrices of each CRE loop sub-cluster as (**a**, **b**), in G1 cells after triptolide treatment. Bin size: 10kb.

**Extended Data Fig. 7.**
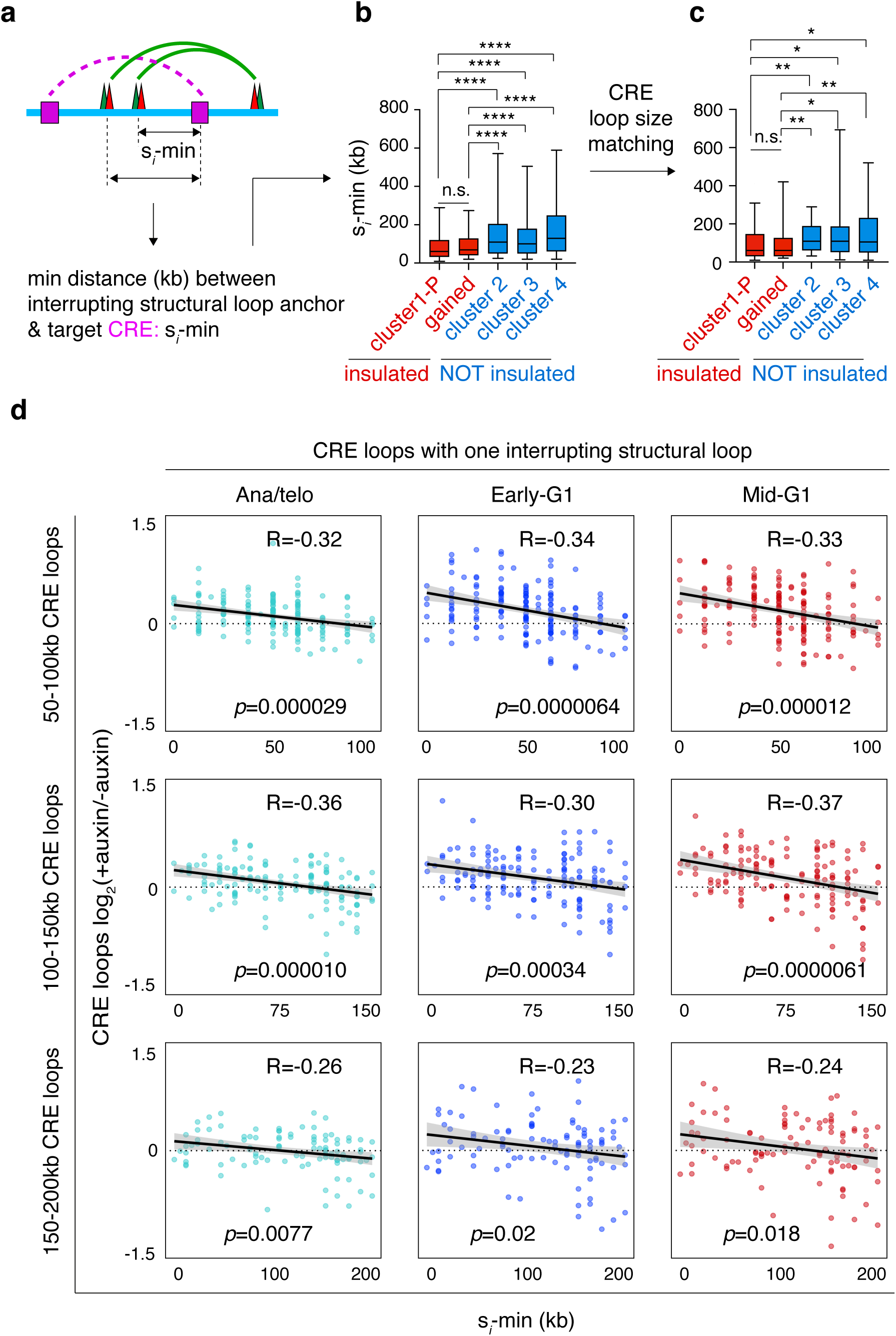
Relative genomic position of structural loops determines their ability to disrupt CRE loops. **a**, Definition of s*_i_*-min. **b**, Boxplots showing significantly lower s*_i_*-min in the insulated cluster1-P and newly gained CRE loops in comparison to the less insulated cluster2, 3 and 4 CRE loops. For all boxplots, central lines denote medians; box limits denote 25th–75th percentile; whiskers denote 5th–95th percentile. * *P* < 0.05, ** *P* < 0.01, *** *P* <0.001 and **** *P* <0.0001. Two-sided Mann-Whitney U test. **c**, Boxplots showing the same comparison as (**b**) after CRE loop size-matching. For all boxplots, central lines denote medians; box limits denote 25th–75th percentile; whiskers denote 5th–95th percentile. * *P* < 0.05, ** *P* < 0.01, *** *P* <0.001 and **** *P* <0.0001. Two-sided Mann-Whitney U test. **d**, Scatter plots showing the significant negative correlation across all post-mitotic cell cycle stages between insulation strength (log_2_ fold change +auxin/-auxin) and s*_i_*-min for CRE loops with a single interrupting structural loop. Top panel: sizes of CRE loops were limited to 50-100kb. Middle panel: sizes of CRE loops were limited to 100-150kb. Bottom panel: CRE loop sizes were limited to 150-200kb. The correlation coefficient and *p* values were computed via the “cor.test” function in R.

**Extended Data Fig. 8.**
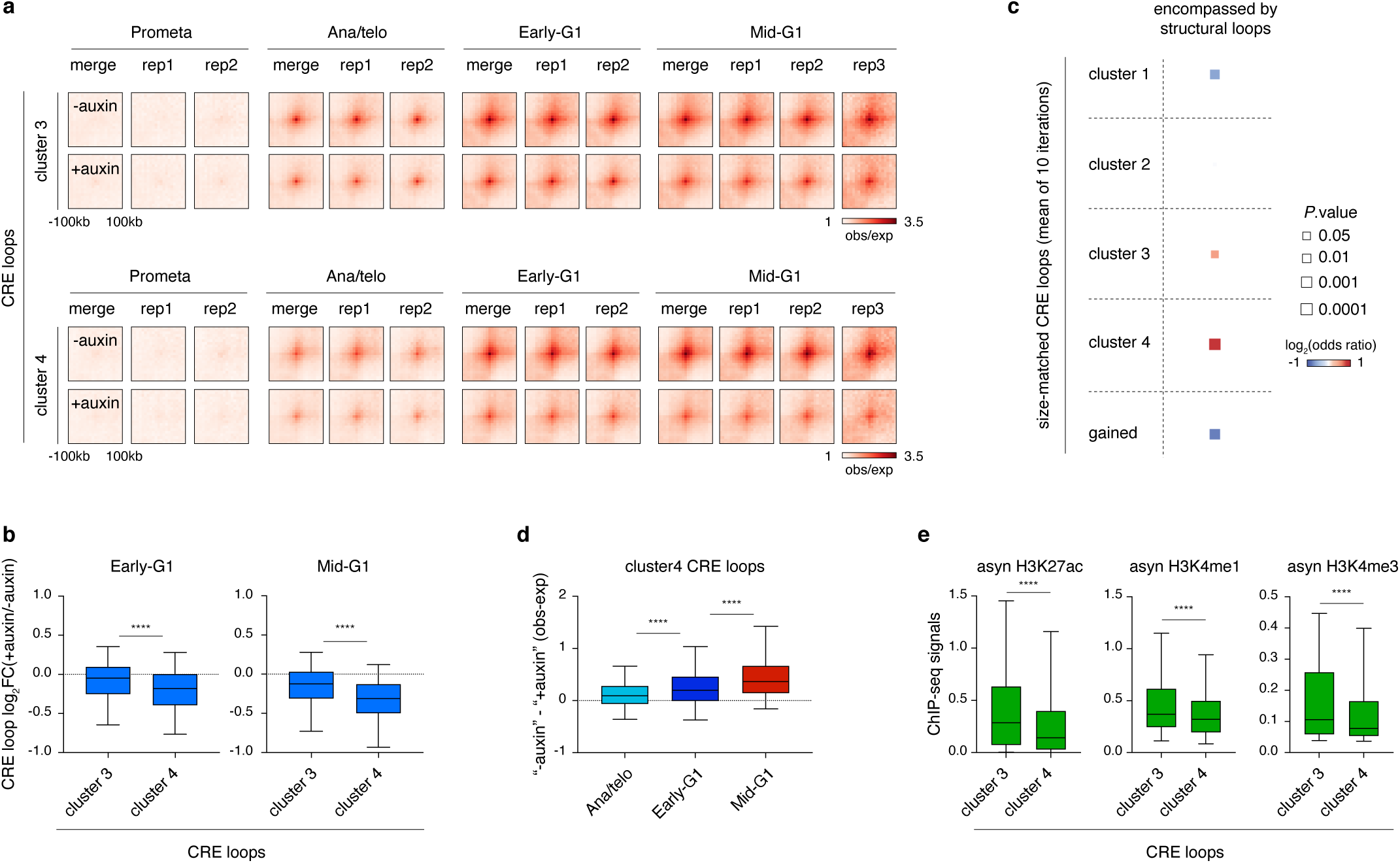
Structural loops support the formation of interactions especially among weak CREs. **a**, APA plots of the cluster3 and cluster4 CRE loops with or without auxin at all tested time points. Bin size: 10kb. Plots for the replicate-merged as well as each individual replicates are shown. **b**, Box plots showing the log_2_ fold change of cluster3 and cluster4 CRE loop strength after auxin treatment in early- and mid-G1 phase. For all boxplots, central lines denote medians; box limits denote 25th–75th percentile; whiskers denote 5th–95th percentile. **** *P* <0.0001. Two-sided Mann-Whitney U test. **c**, Enrichment analysis of each CRE loop cluster encompassed by structural loops. Colors of the squares indicate log_2_ transformed odds ratio (Fisher’s exact test). Sizes of squares indicate the significance of enrichment (*p* values of Fisher’s exact test). **d**, Box plots showing the progressively strengthened support (loop strength “-auxin” – “+auxin”, obs/exp) by structural loops of cluster4 CRE loops. For all boxplots, central lines denote medians; box limits denote 25th–75th percentile; whiskers denote 5th–95th percentile. **** *P* <0.0001. Two-sided paired Wilcoxon signed-rank test. **e**, Box plots showing the ChIP-seq signals of indicated histone marks at anchors of cluster3 or cluster4 CRE loops. For all boxplots, central lines denote medians; box limits denote 25th–75th percentile; whiskers denote 5th–95th percentile. **** *P* <0.0001. Two-sided Mann-Whitney U test.

**Extended Data Figure 9.**
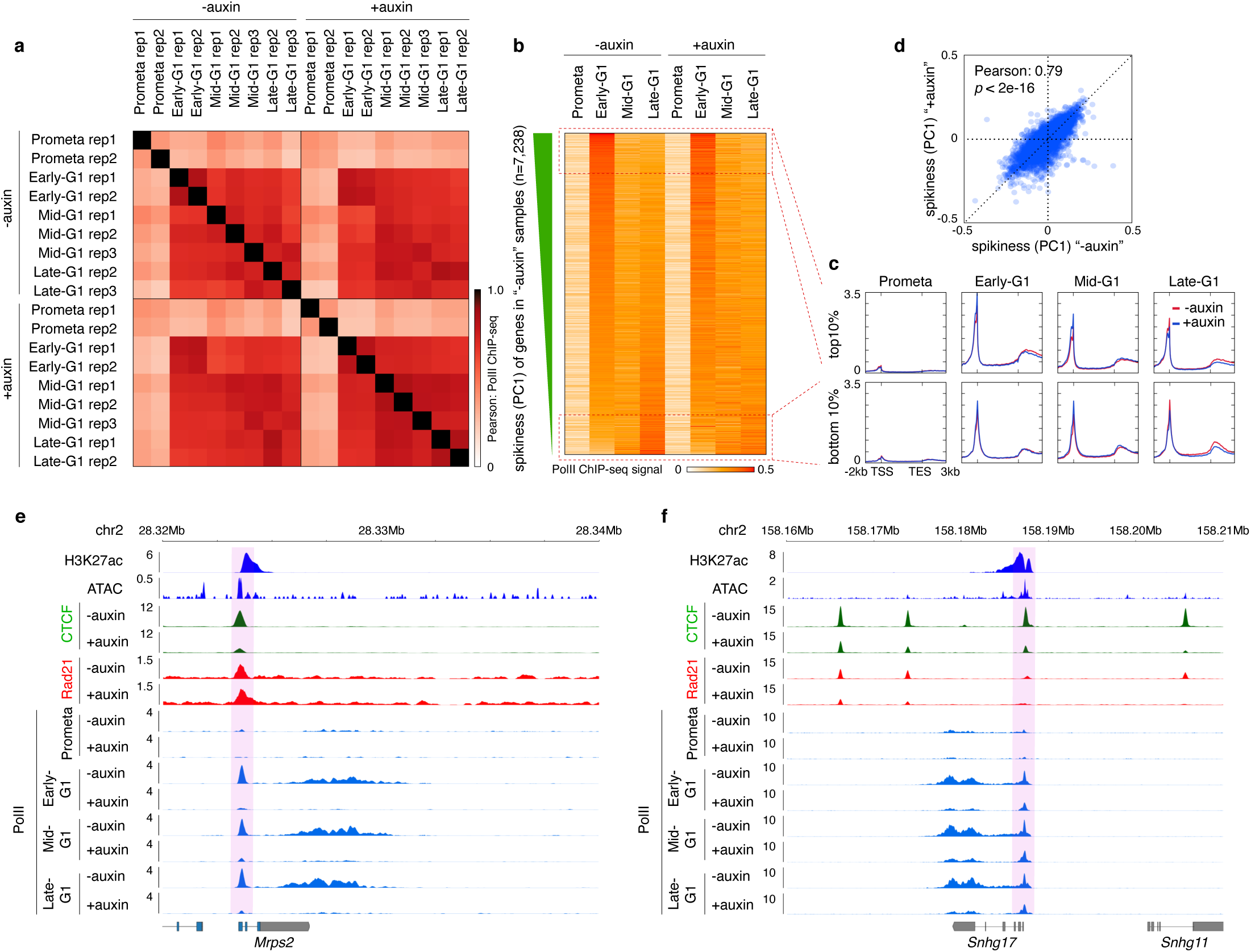
Characterization of post-mitotic gene reactivation after CTCF depletion. **a**, Pearson correlations of PolII ChIP-seq signals across biological replicates. **b**, Heatmap of post-mitotic transcriptional “spikiness” of genes in untreated and auxin treated samples. Active genes were ranked in a descending order of the PC1 of untreated samples. **c**, Meta-region plots (scaled from TSS to TES) showing the PolII ChIP-seq signals of the 10% most spiking and least spiking genes across cell cycle stages with or without CTCF. **d**, Scatter plot showing the positive correlation between PC1 values of untreated and auxin treated sample, confirming that post-mitotic spiking is largely preserved after CTCF depletion. **e**, Browser tracks of the *Mrps2* locus showing dramatically reduced PolII ChIP-seq signals after CTCF depletion. Note a strong CTCF peak located at the TSS that was diminished upon auxin treatment. However, the Rad21 signal was minimally perturbed after CTCF depletion at this site, suggesting that cohesin is loaded at this region. **f**, Similar to (**e**), showing the PolII signal reduction at *Snhg17* locus.

**Extended Data Figure 10.**
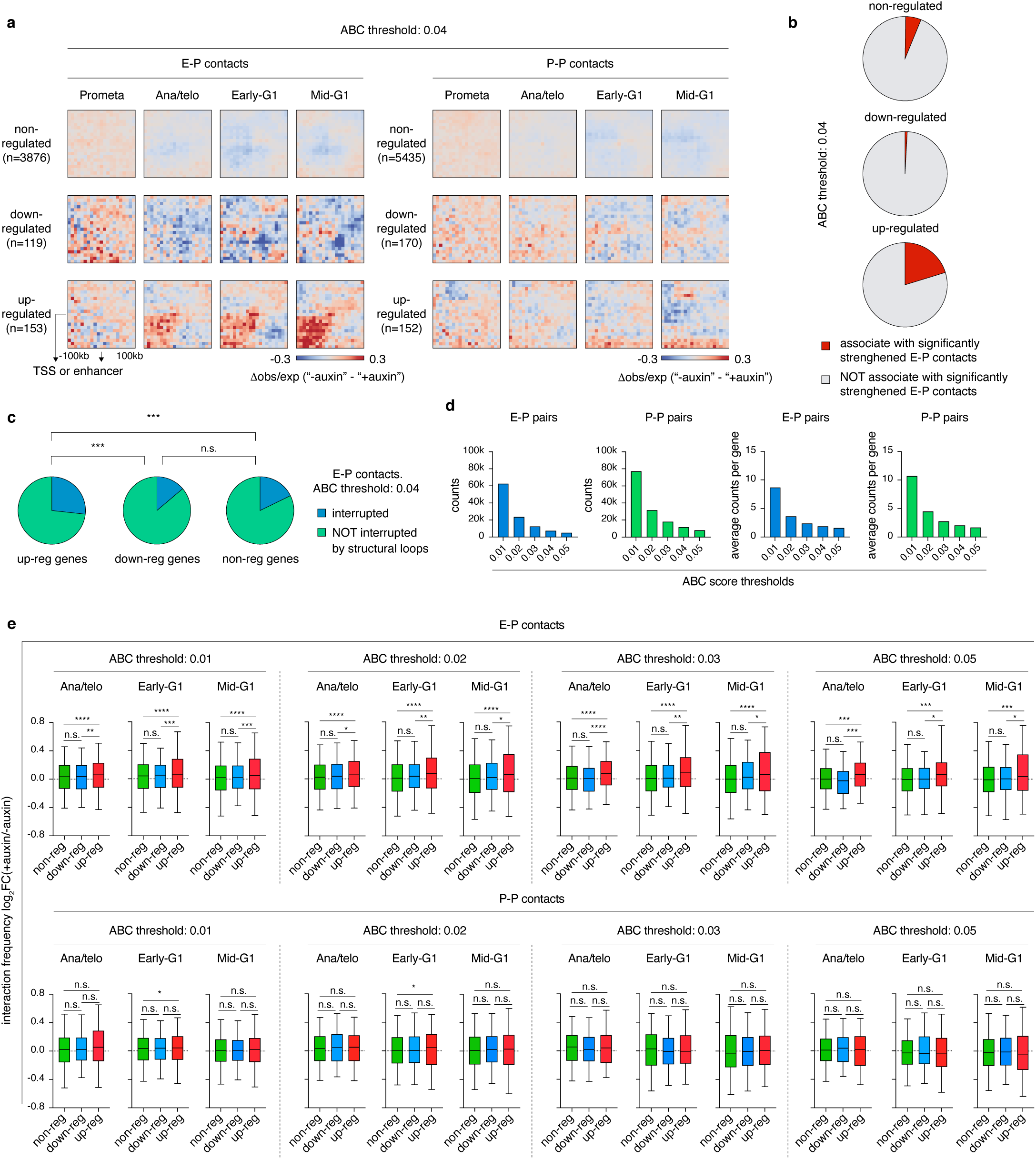
CTCF loss-actuated post-mitotic gene up-regulation is linked to elevated enhancer-promoter interactions. **a**, Left: Pile-up Hi-C matrices showing the changes of E-P interactions associated with non-regulated, down-regulated or up-regulated genes after CTCF depletion. Bin size: 10kb. Note that increase of E-P interactions for up-regulated genes was observed as early as in ana/telophase. Right: Similar to left with pile-up analysis showing changes of P-P interactions. Bin size: 10kb. **b**, Pie charts showing the fraction of up-, down- or non-regulated genes that are associated with significantly strengthened E-P contacts. **c**, Pie charts showing the fraction of structural loop-interrupted E-P pairs associated with non-regulated, down-regulated or up-regulated genes when ABC cutoff equals 0.04. *** *P* <0.001. *P* values were calculated using Fisher’s exact test. **d**, Numbers of total and per gene confident E-P pairs or P-P pairs with ABC score cutoffs set to 0.01, 0.02, 0.03, 0.04 or 0.05. **e**, Similar to Figure 4e and f, with Boxplots showing the log_2_ fold change of interactions between E-P pairs or P-P pairs associated with non-regulated, down-regulated and up-regulated respectively with ABC scores set to 0.01, 0.02, 0.03 or 0.05. For all boxplots, central lines denote medians; box limits denote 25th–75th percentile; whiskers denote 5th–95th percentile. * *P* < 0.05, ** *P* < 0.01, *** *P* <0.001 and **** *P* <0.0001. Two-sided Mann-Whitney U test.

**Extended Data Figure 11.**
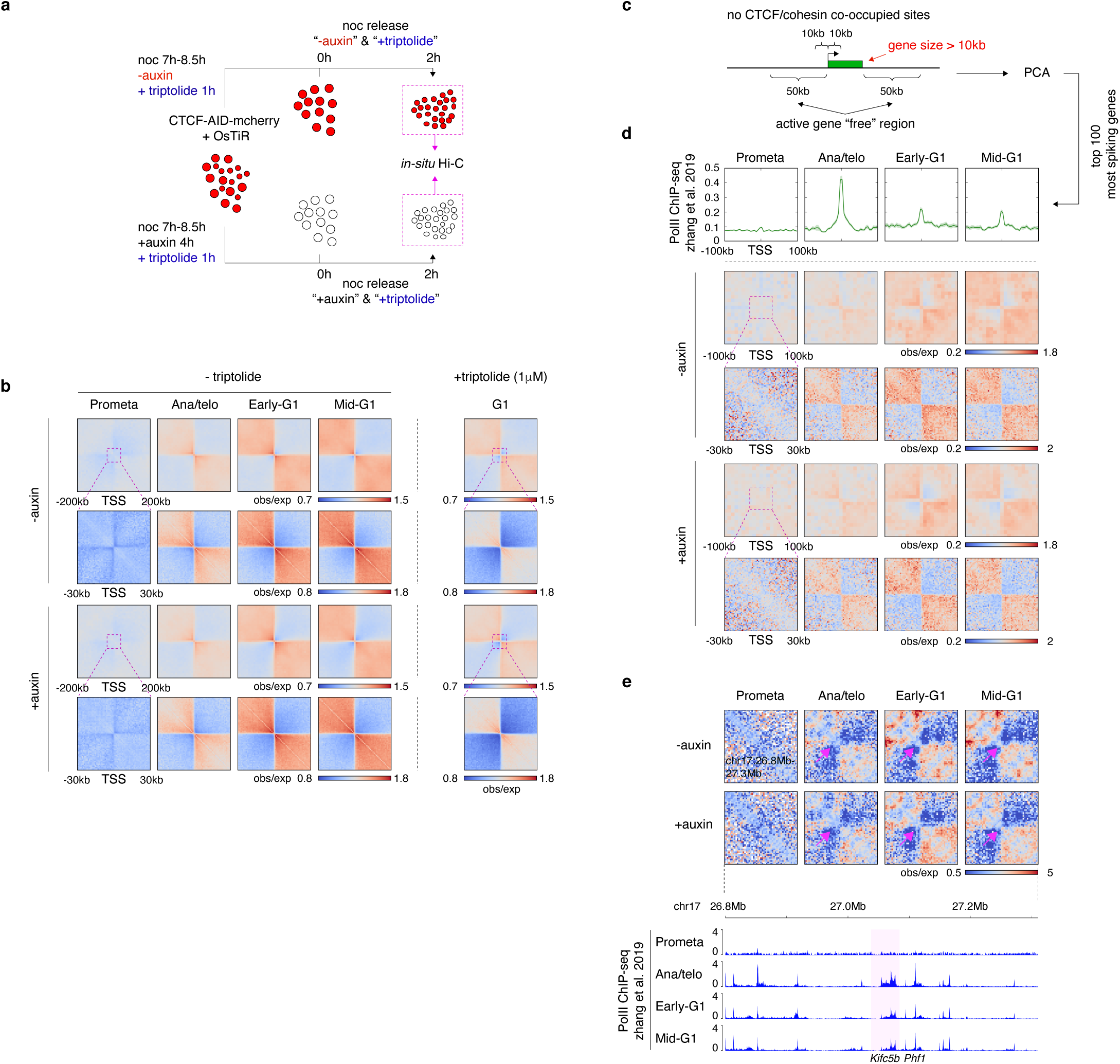
Relationship between TSS PolII binding and local insulation. **a**, Outline of experimental flow involving transcriptional inhibition and CTCF depletion during the mitosis-to-G1 phase transition. **b**, Pile-up Hi-C matrices showing the insulation with or without triptolide treatment at all active TSS identified previously^1^. Insulation plots were generated across cell cycle stages in both untreated control and auxin treated cells. Plots with bin size of 10kb and 1kb (zoom-in view) are shown. **c**, Strategy of target gene selection (see methods) to find genes with PolII spiking at the TSS. **d**, Upper panel: Meta-region plots showing the PolII ChIP-seq profiles (from parental G1E-ER4 cells) of the top 100 most spiking TSS chosen from (**c**). Plots were centered on TSS. Lower panel: Pile-up Hi-C matrices showing the insulation at the top 100 most spiking TSS chosen from (**c**). Plots with bin size of 10kb and 1kb (zoomed-in view) were plotted. **e**, Upper panel: KR balanced Hi-C contact matrices showing the progressive insulation gain at the *Kifc5b* and *Phf1* loci in untreated control and auxin treated samples. Insulation is indicated by purple arrows. Bin size: 10kb. Lower panel: Genome browser tracks showing PolII occupancy corresponding to genomic the region in upper panel across cell cycle stages in parental cells.

**Extended Data Figure 12.**
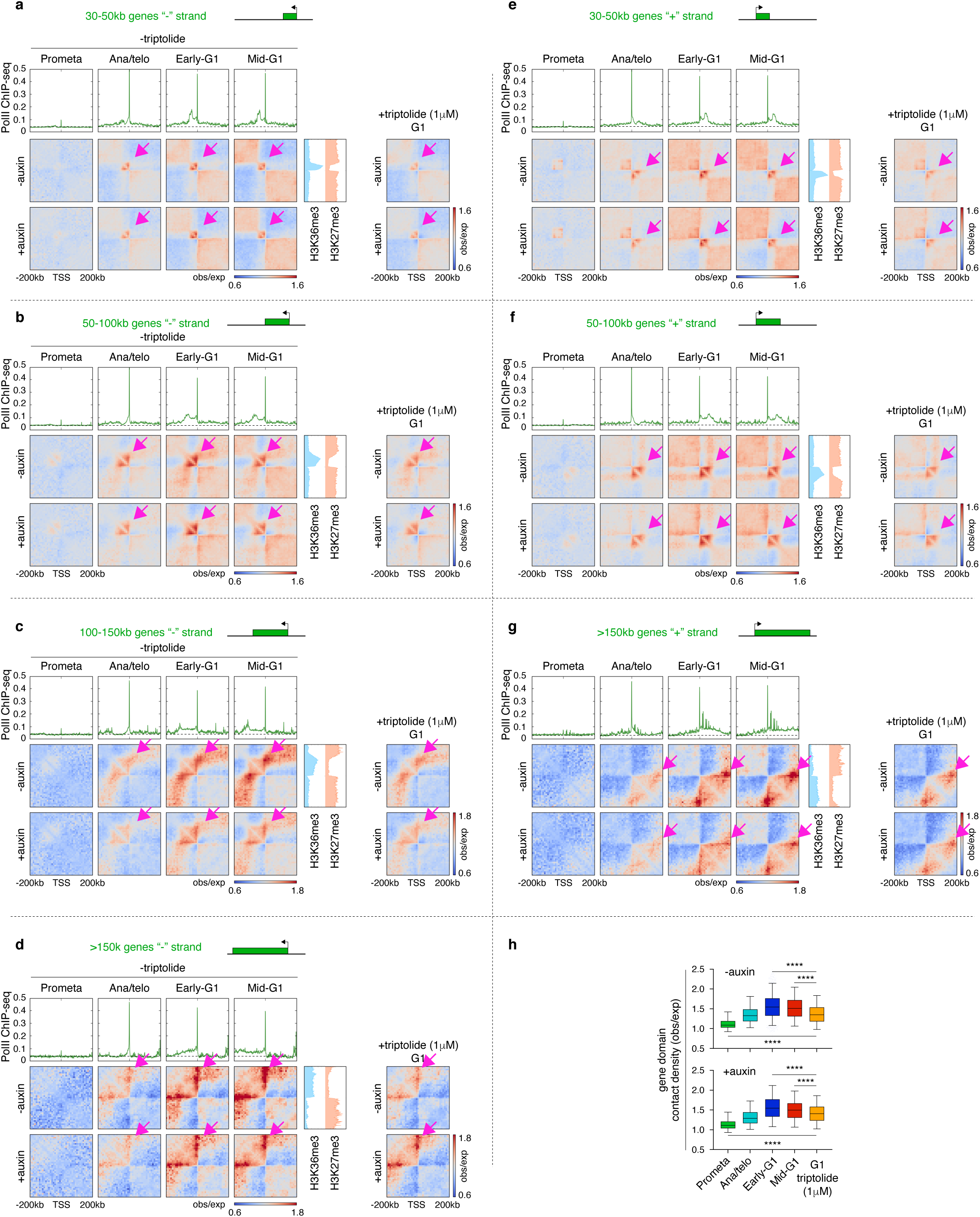
Partial uncoupling of gene domain formation from active transcription after mitosis. **a**, Upper panels: Meta-region plots showing the parental PolII ChIP-seq profile of all minus strand genes between 30kb and 50kb across all cell cycle stages. Plots were centered on TSS. Bottom panels: Pile-up Hi-C matrices showing the progressive reformation of gene domains corresponding to the upper panel in samples with or without triptolide treatment. Corresponding samples after auxin treatment are also shown. Plots are centered around TSS. Gene domains are indicated by purple arrows. Meta-region plots of showing H3K36me3 and H3K27me3 ChIP-seq are sh. **b**-**g**, Similar to (**a**) showing PolII elongation and reformation of gene domains with indicated gene size range and strandedness in samples with or without triptolide treatment. Corresponding samples with auxin treatment are also shown. **h**, Box plots showing the strength of gene domains in post-mitotic samples without triptolide treatment as well as G1 samples with triptolide treatment. For all boxplots, central lines denote medians; box limits denote 25th–75th percentile; whiskers denote 5th–95th percentile. **** *P* <0.0001. Two-sided Paired Wilcoxon signed rank test.

Supplementary table 1: Hi-C data processing statistics

Supplementary table 2: rGMAP boundary calls

Supplementary table 3: Hiccups loop calls

Supplementary table 4: Active genes

